# A cascade of sulfur transferases delivers sulfur to the sulfur-oxidizing heterodisulfide reductase-like complex

**DOI:** 10.1101/2023.12.18.572138

**Authors:** Tomohisa Sebastian Tanabe, Elena Bach, Giulia D’Ermo, Marc Gregor Mohr, Natalie Hager, Niklas Pfeiffer, Marianne Guiral, Christiane Dahl

## Abstract

A heterodisulfide reductase-like complex (sHdr) and novel lipoate-binding proteins (LbpAs) are central players of a wide-spread pathway of dissimilatory sulfur oxidation. Bioinformatic analysis demonstrate that the cytoplasmic sHdr-LbpA systems are always accompanied by sets of sulfur transferases (DsrE proteins, TusA, rhodaneses). The exact composition of these sets may vary depending on the organism and sHdr system type. To enable generalizations, we studied model sulfur oxidizers from distant bacterial phyla, i.e. Aquificota and Pseudomonadota. DsrE3C of the chemoorganotrophic Alphaproteobacterium *Hyphomicrobium denitrificans* and DsrE3B from the Gammaproteobacteria *Thioalkalivibrio* sp. K90mix, an obligate chemolithotroph, and *Thiorhodospira sibirica*, an obligate photolithotroph, are homotrimers that donate sulfur to TusA. Additionally, the hyphomicrobial rhodanese-like protein Rhd442 exchanges sulfur with both TusA and DsrE3C. The latter is essential for sulfur oxidation in *Hm. denitrificans*. TusA from *Aquifex aeolicus* (AqTusA) interacts physiologically with AqDsrE, AqLbpA and AqsHdr proteins. This is particularly significant as it establishes a direct link between sulfur transferases and the sHdr-LbpA complex that oxidizes sulfane sulfur to sulfite. *In vivo,* it is unlikely that there is a strict unidirectional transfer between the sulfur-binding enzymes studied. Rather, the sulfur transferases form a network, each with a pool of bound sulfur. Sulfur flux can then be shifted in one direction or the other depending on metabolic requirements. A single pair of sulfur-binding proteins with a preferred transfer direction, such as a DsrE3-type protein towards TusA, may be sufficient to push sulfur into the sink where it is further metabolized or needed.

**SIGNIFICANCE STATEMENT:** A network of bacterial sulfur transferases is uncovered and characterized that ultimately delivers sulfur to a complex cytoplasmic sulfur-oxidizing metalloenzyme, sHdr, that resembles heterodisulfide reductase from methanogenic archaea and interacts with lipoate-binding proteins. Similar sets of sulfur transferases occur in phylogenetically distant bacteria, underscoring the fundamental importance of the work.

## 1. INTRODUCTION

In the cytoplasm of prokaryotes, as in most (if not all) other biological contexts, reduced sulfur is normally handled in a protein-bound state due to its reactivity.^(1–5)^ Often, multiple sulfur transferases form a network to direct the sulfur to the correct metabolic pipeline or its target molecule. In these networks, individual sulfur-trafficking proteins may provide sulfur to multiple target reactions/proteins. Such networks are not only important for the biosynthesis of sulfur-containing cellular components but also essential in many Bacteria and Archaea, which carry out dissimilatory sulfur oxidation with energy conservation via respiratory or photosynthetic electron transport.^(6,7)^ Here, the enzymatic production of persulfide sulfur, the successive transfer of sulfur as a persulfide between multiple proteins, and the oxidation of sulfane sulfur in protein-bound form are all crucial steps. Rhodaneses, TusA and DsrE-like sulfur transferases are central and common elements in these processes. They have an established role in the rDsr pathway of sulfur oxidation where they work together to transfer sulfur as a cysteine persulfide to DsrC, which ultimately presents the sulfur to the oxidizing enzymatic unit, dissimilatory sulfite reductase, DsrAB.^(6,8,9)^ In the model organism *Allochromatium vinosum,* sulfur atoms are successively transferred from the rhodanese Rhd_2599 to TusA,^(8)^ then to a conserved cysteine residue of DsrE, the active site subunit of the heterohexameric DsrE_2_F_2_H_2_ complex,^(10)^ and from there to DsrC.^(9)^ The function of the membrane-bound AvDsrE2A protein, another sulfur transferase involved in this enzymatic relay, remains unclear.^(8)^

Recently, the sulfur-oxidizing heterodisulfide reductase-like sHdr complex and novel lipoate-binding proteins (LbpAs) have been identified as central players of an additional widespread sulfur oxidation pathway.^(11–13)^ It exists in a significant group of organisms that comprise the volatile organic sulfur compound degrader *Hyphomicrobium denitrificans*, along with many chemo- and photolithoautotrophic bacteria and archaea, that include several environmentally relevant sulfur oxidizers such as *Acidithiobacillus sp.*, *Thioalkalivibrio* or *Sulfobacillus* species and the hyperthermophile *Aquifex aeolicus*. Two types of gene *shdr* clusters can be differentiated (Figure 1): Type I with an *shdrC1B1AHC2B2* arrangement and type II with an *shdrC1B1AHB3-etfAB-emo* arrangement.^(11,14)^ While the importance of the type I cluster for sulfur dissimilation has been proven,^(11–13)^ less is known for type II, although prokaryotes encoding the type II system are known sulfur oxidizers.^(14)^ Irrespective of the type of gene cluster, the *shdr-lbpA* genes are conspicuously often associated with genes for accessory sulfurtransferases similar to but not identical with those that fuel the rDsr pathway (Figure 1).^(11,12)^

**FIGURE 1.**
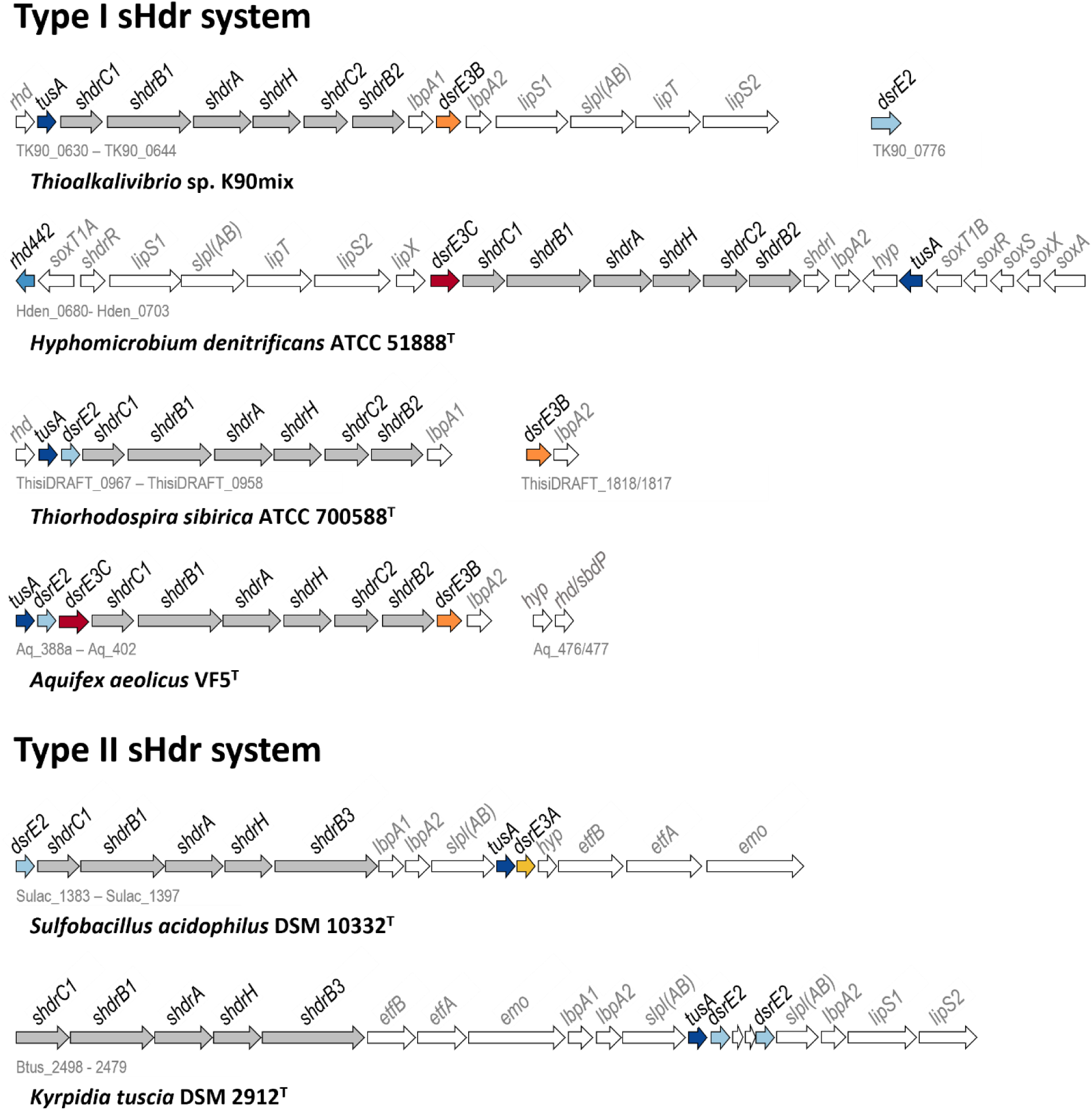
Representative *shdr* gene clusters in sulfur oxidizers. The KEGG/NCBI locus tag identifiers for the first and last genes are shown below each cluster. Genes for TusA, DsrE2, DsrE3A, DsrE3B and DsrE3C are indicated in dark blue, light blue, yellow, orange and dark red, respectively. Genes for probable components of the sulfur-oxidizing heterodisulfide reductase-like (sHdr) complex are shaded in gray. EMO, ETF:(methyl)menaquinone oxidoreductase; Etf, electron transfer protein. EtfAB and EMO have been proposed to direct electrons stemming from sulfane sulfur oxidation to menaquinone.^(17)^ LbpA, lipoate-binding protein; sLpl(AB), lipoate:protein ligase; LipS1/S2, lipoyl synthase; LipT, FAD-binding NAD(P)H-dependent oxidoreductase possibly delivering electrons for the LipS1/S2-catalyzed sulfur insertion step ^(13,17)^; Rhd, rhodanese; SbdP, sulfur-binding-donating protein.

Biochemical information regarding sulfur transferases associated with sHdr is extremely limited. The only sHdr-associated sulfur transferases studied thus far come from the archaeon *Metallosphaera cuprina*. Its DsrE3A protein has been biochemically characterized as a thiosulfonate transferase.^(15)^ Bound thiosulfate is transferred from DsrE3A to TusA but not vice versa^(15)^ implying that DsrE3A functions as a thiosulfate donor to TusA *in vivo*. The DsrE2A-type sulfur transferases, which are encoded in certain *shdr* gene clusters (Figure 1), have been proposed as potential membrane anchors for the sHdr-like complex.^(16)^

Here we set out to shed more light on sulfur transfer to the sHdr system. Initially, we analyse the potential correlation between the prevalence of *shdr* and sulfur transferases genes. To enable generalizations, we concentrate on the type I sHdr system in bacteria and our model organisms stem from two distant phyla: the Aquificota (*Aquifex aeolicus*) and the Pseudomonadota, which are represented by sulfur oxidizers from two different classes, the Alphaproteobacteria (*Hyphomicrobium denitrificans*) and the Gammaproteobacteria (*Thiorhodospira sibirica* and *Thioalkalivibrio* sp. K90mix). From these organisms, we investigate the proteins Rhd442, TusA, DsrE3B and DsrE3C to determine whether and how they mobilize and transfer sulfur. In the genetically accessible *Hm. denitrificans*, we gather information on the importance of DsrE3C *in vivo*.

## 2 RESULTS

### 2.1 Distribution of the sHdr system

Clusters of genes encoding the sHdr pathway for sulfane sulfur oxidation in the cytoplasm fall into two distinct categories^(11,14,17)^ (Figs. 1 and 2). The type I and type II sHdr systems share several core proteins, namely the Fe/S-flavoprotein sHdrA, the electron carrier protein sHdrC1, that binds two cubane [4Fe-4S] clusters and the proposed catalytic subunit sHdrB1 that probably coordinates two noncubane Fe/S clusters.^(18)^ sHdrC2 is another ferredoxin-like electron carrier. sHdrB2 has the potential to bind two classical noncubane Fe/S clusters and probably acts as a disulfide reductase.^(18)^ Organisms with type II sHdr systems encode a protein that we term sHdrB3. This is a fusion of sHdrC2 and sHdrB2, albeit it can bind only one noncubane Fe/S cluster.^(17)^ Electron transfer protein EtfAB and ETF:(methyl)menaquinone oxidoreductase EMO are encoded within type II *shdr* clusters and have been proposed to direct electrons stemming from sulfane sulfur oxidation to menaquinone.^(17)^

**FIGURE 2.**
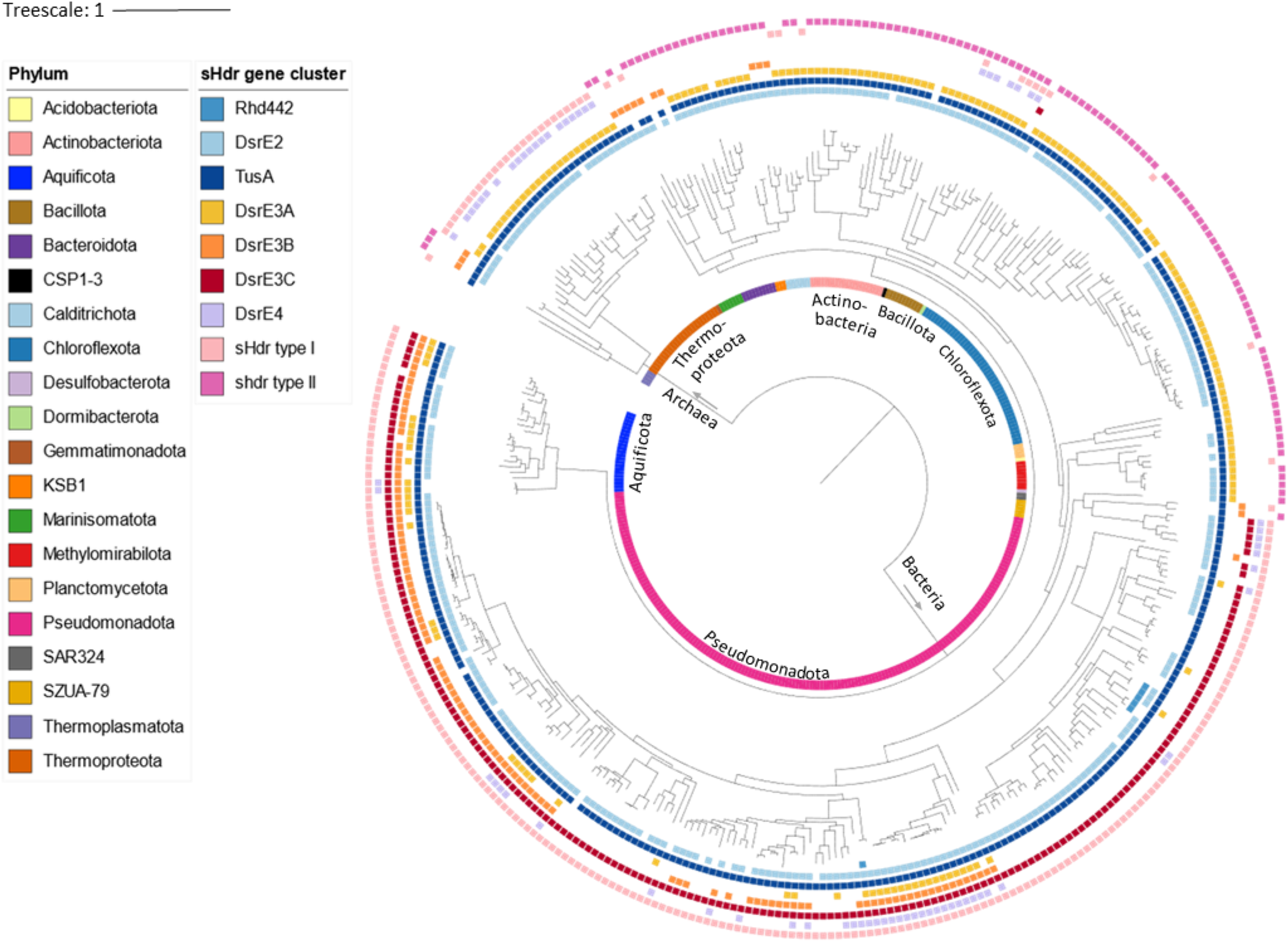
Taxonomic distribution of type I or type II sHdr systems in sulfur-oxidizing prokaryotes. The distribution of TusA, Rhd442 and DsrE-type sulfur transferases is also visualized. The data underlying the figure is provided in Table S1.

Of the entire GTDB representative genome collection (release R207), 397 assemblies from 20 phyla contain core *shdr* genes. For 353 of these assemblies, the concatenated sequences for 16 ribosomal proteins could be used as phylogenetic markers to compute a species tree,^(19,20)^ that served as the basis for mapping the distribution of type I and II sHdr systems using HMSS2^(21)^ (Figure 2, Table S1). Among the Bacteria, the Pseudomonadota, the Aquificota, and the phylum SZUA-79 have exclusively the type I system. Among the Archaea, the type I *shdr* gene set occurs in the Thermoproteota, a phylum harboring well-established sulfur oxidizers like *Sulfolobus sp.* and *Acidianus* sp. The type II sHdr system is found in 130 assemblies from one archaeal and 12 bacterial phyla. The majority of the respective bacterial assemblies belong to the Chloroflexota, Actinobacteriota and Bacillota. In some bacterial phyla, there are species that have either type I or type II sHdr systems (e.g. Marinisomatota or Chloroflexota). All sHdr-containing members of the Bacillota contain the type II system. In addition, four *Sulfobacillus* species, which are well-established sulfur oxidizers,^(14,22–24)^ and one further member of the order Sulfobacillales bear the genetic capacity for both sHdr systems (Figure 2, Table S1).

### 2.2 Distribution of sHdr-associated sulfurtransferases

Here, we intended to further illuminate the general association of *shdr* gene clusters with genes for different sulfurtransferases^(2,11,15,17)^ (TusA, DsrE-type sulfur transferases, rhodanese Rhd442). In order to do so, we first needed to clearly describe and validate the various classes of proteins using sequence similarity networks (SSN). Clusters in SSNs reflect phylogenetic clades.^(25)^

TusA family proteins are a central hub in sulfur transfer during various anabolic and catabolic processes.^(1)^ In *Escherichia coli*, the three homologous TusA-family proteins, TusA, YedF and YeeD have distinct functions and cannot substitute for each other.^(26)^ Besides contributing to tRNA thiolation,^(27)^ TusA mediates sulfur transfer for molybdenum cofactor biosynthesis and affects iron-sulfur cluster assembly as well as the activity of major regulatory proteins.^(26,28,29)^ YeeD is a component of thiosulfate uptake for sulfur assimilation.^(30)^ YedF in some way affects flagella formation and motility.^(31)^ We retrieved all sequences clustering with these three proteins from Uniprot and subjected them to a SSN analysis together with TusA-like proteins that are genetically linked with sulfur-oxidation systems. Indeed, TusA, YedF, and YeeD each form distinct clusters of similar sequences (Figure 3a). In addition, three further clusters are clearly discernible. Two of them consist of TusA-like proteins linked with rDsr systems and a third comprises TusAs linked with sHdr and rDsr-systems. In summary, TusAs genetically associated with sulfur oxidation systems can be confidently distinguished from classical TusA, YedF and YeeD and form distinct phylogenetic clades (Figure 3a).

**FIGURE 3.**
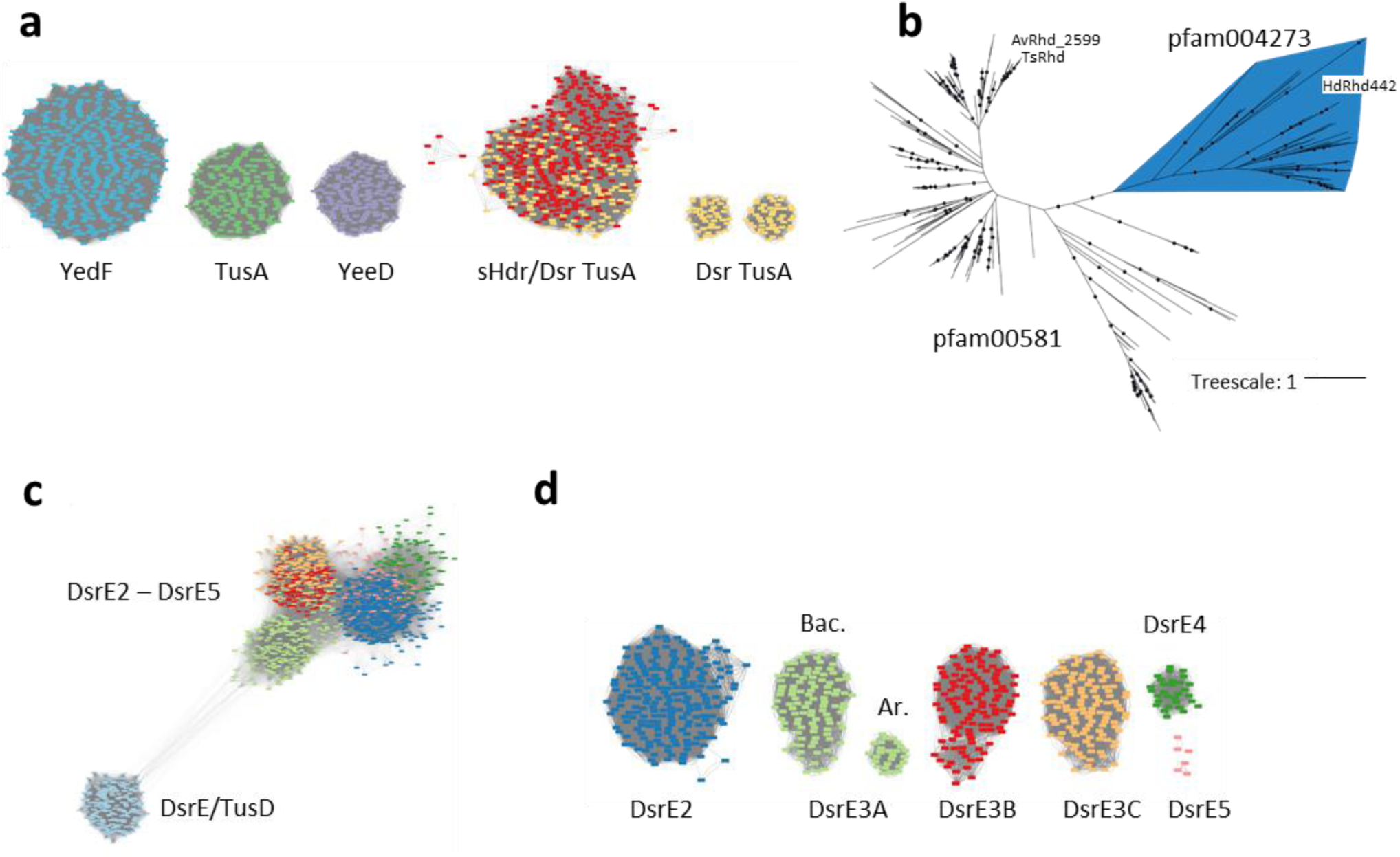
Sequence similarity network (SSN) and phylogenetic analyses for sulfur transferases relevant for dissimilatory sulfur oxidation. (a) SSN for the TusA family (pfam01206) members clustering at 50 % identity with TusA, YedF and YeeD from *E. coli* and TusA-like proteins that are genetically linked with sHdr and/or Dsr sulfur-oxidation systems. Connections below a threshold score of 100 were removed. (b) Maximum likelihood tree for ∼100-aa single domain rhodaneses that are encoded in synteny with DsrE2 and TusA or that are related to Rhd442 from *Hm. denitrificans* X^T^. Bootstrap values exceeding 95% are indicated by dots. (c) SSN for all DsrE-type sequences from the representative genomes of GTDB (release R207). (d) SSN for all DsrE homologs associated with rDsr and/or sHdr systems, excluding the divergent TusD/DsrE proteins. Connections below a threshold score of 135 were removed. Bac., Bacteria; Ar, Archaea.

Rhodaneses were originally identified and named on the basis of their ability to catalyze the transfer of a sulfane sulfur atom from thiosulfate to cyanide yielding SCN^-^ (rhodanide, thiocyanate) as the product.^(32)^ Rhodaneses and rhodanese-like proteins are very widespread^(33)^ and therefore we did not consider it useful to create an SSN analysis spanning all prokaryotes, but instead limited ourselves to sulfur oxidizers with rDsr and/or sHdr systems. Within a first group of these, the corresponding gene is present in a syntenic cluster with genes for DsrE2 and TusA (e.g. in *Ts. sibirica*, Figure 1). A second group contains proteins related to rhodanese Rhd442 encoded in the vicinity of the *shdr* gene cluster in *Hm. denitrificans* X^T^ (Figure 1). By establishing a phylogenetic tree (Figure 3b), it became apparent that the DsrE2-TusA associated rhodaneses belong to a different clade than the Rhd442-related proteins (Figure 3b). The former belong to a protein family (pfam00581) that also encompasses Rhd_2599 from *Ac. vinosum*^(8)^ and SbdP from *Aq. aeolicus*.^(34)^ While the first is part of the relay delivering sulfur to the rDsr pathway of sulfur oxidation,^(8)^ SbdP can load long sulfur chains and interacts with sulfur reductase and sulfur oxygenase reductase, i.e. key enzymes of sulfur energy metabolism.^(34)^ Rhd442 belongs to a separate protein family, pfam004273 (DUF442) that forms a monophyletic group in the rhodanese tree presented Figure 3b.

Originally, the DsrE family has been categorized into five well distinguishable phylogenetic groups, DsrE, DsrE2, DsrE3 (with subgroups DsrE3A, DsrE3B and DsrE3C), DsrE4 and DsrE5.^(15,16,35)^ Here, we clustered all DsrE-type sequences from the representative genomes of GTDB (release R207) by sequence similarity network (SSN) analysis. In a first approach, the proteins were not filtered for association with dissimilatory sulfur oxidation. Classical TusD/DsrE were clearly distinguishable and only distantly related to the sulfurtransferases from the other subclasses (Figure 3c). In a second approach, an SSN was calculated with all DsrE homologs associated with rDsr and/or sHdr systems, excluding the divergent TusD/DsrE proteins (Figure 3d). For DsrE2, DsrE3 and DsrE4 robust clades mirroring the original DsrE groups and subgroups were retrieved.^(15)^ DsrE3A formed two robust phylogenetic clades, containing archaeal and bacterial sequences, respectively (Figure 3d). DsrE5 proteins present an exception as they do not cluster as a coherent group due to high sequence dissimilarity. While functional predictions for this group are not available, members of the DsrE4 group are proposed to play a role in detoxification of reactive sulfur species.^(15)^ As already pointed out, archaeal DsrE3A is an established thiosulfonate carrier whereas DsrE3B and DsrE3C have not been studied on a biochemical level yet. Although a genetic association indicates a function in the sHdr system, the mechanism and specific roles of DsrE3B and DsrE3C have not been described so far.

Once the sulfurtransferases could be confidently categorized, their co-occurrence in sHdr-containing organisms was studied using HMSS2.^(21)^ In addition, they were mapped onto the phylogenetic tree shown in Figure 2. Of all 397 studied assemblies, 387 encode at least one TusA and 302 of the *shdr* clusters are directly linked to *tusA* genes. The *dsrE2* gene is present in 345 (87%) assemblies and in 199 (50.1%) of the *shdr* gene clusters, making it the second most frequently occurring gene. The *dsrE3A* (200), *dsrE3B* (132), and *dsrE3C* (223) genes are less prevalent. However, when these genes are present, 70%, 90% and 52%, respectively, are located in the immediate vicinity of *shdr* core genes. The rhodanese-encoding gene *rhd442* is present in only three *shdr* gene clusters from the family *Hyphomicrobiaceae* (Alphaproteobacteria).

The distribution of DsrE-type sulfurtransferases appears to relate to the type of sHdr system (Figure 2). Archaea with type I sHdr system particularly often contain *shdr* genes linked with *tusA*, *dsrE3A* and *dsrE4*. This is different in Bacteria with type I sHdr system, where a *tusA*/*dsrE3C* combination occurs in the same genome at a notably high frequency. Other DsrE-type sulfur transferases may be present but are less abundant. Some members of the Pseudomonadota and Aquificota simultaneously encode DsrE3A and DsrE3B. The highest number of different sulfurtransferases is found in genomes from the gammaproteobacterial orders Acidithiobacillales, Acidiferrobacterales and Ectothiorhodospirillales (here in the genus *Thioalkalivibrio* and in the family Acidihalobacteraceae). These organisms have the genetic potential for DsrE2, DsrE3A, DsrE3B, DsrE3C and DsrE4. Type II sHdr systems are predominantly associated with TusA and DsrE3A, while DsrE3B and DsrE3C are only very rarely present (Figure 2). Among the type II-containing archaea of the phylum Thermoplasmatota, the sulfur transferase DsrE3B is the only transferase present and is genetically associated with the *shdr* cluster.

### 2.3 Properties of sHdr associated TusA

The TusA proteins from *Hm. denitrificans* (HdTusA), *Thioalkalivibrio sp.* K90mix (TkTusA), *Ts. sibirica* (TsTusA) and *Aq. aeolicus* (AqTusA), all of which are encoded in type I *shdr* gene clusters, were selected as model proteins for further analysis. All four proteins contain a highly conserved cysteine within an N-terminal CPXP motif, which is characteristic for the active site of TusA proteins (Figure 4a). Leucine and isoleucine have been described at the X position for TusA proteins involved in sulfur oxidation.^(1)^ A second cysteine is present at equivalent positions in AqTusA, HdTusA and *E. coli* TusA (Figure 4a). This cysteine is not involved in sulfur transfer in *E. coli*.^(36)^

**FIGURE 4.**
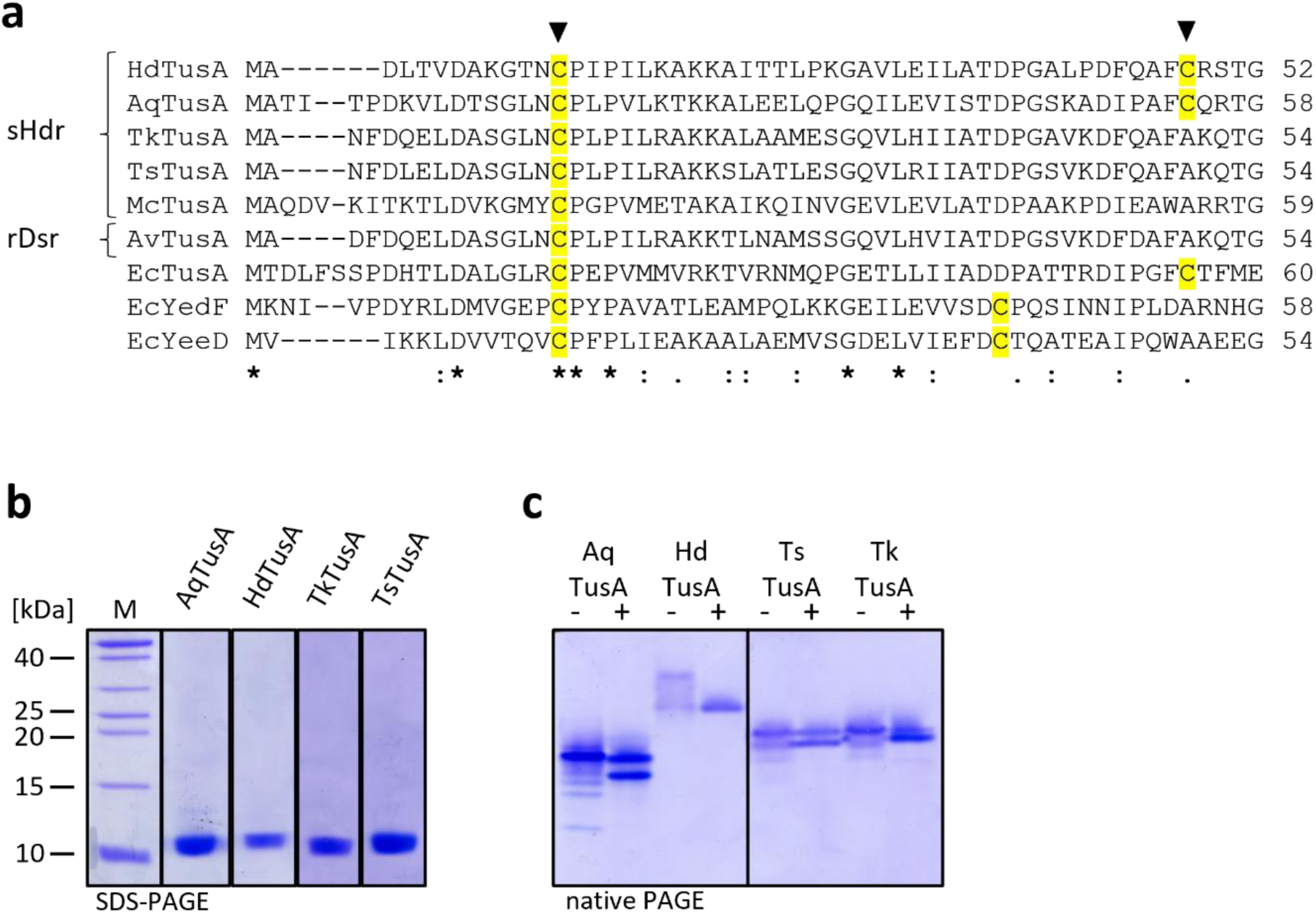
(a) Alignment of a portion of TusA and related proteins from sulfur oxidizers and *E. coli*. Ec, *E. coli* (TusA, b3470, YedF, b1930, YeeD, b2012); Ts, *Ts. sibirica* (ThisiDRAFT_0966); Tk, *Thioalkalivibrio* sp. K90mix (TK90_0631); Av, *Ac. vinosum* (Alvin_2600); Hd, *Hm. denitrificans* (Hden_0698); Aq, *Aq. aeolicus* (Aq_388a); Mc, *Ms. cuprina* (Mcup_0683). Triangles indicate the cysteines that were exchanged to serine in this work. Asterisk, fully conserved residues; colon, conservation between groups of strongly similar properties; dot, conservation between groups of weakly similar properties. (b) 20 % SDS-PAGE of recombinant TusAs HdTusA, TkTusA, TsTusA and AqTusA reduced with DTT. (c) 20 % native PAGE of 3,5 µg recombinant TusAs HdTusA, TkTusA and TsTusA and AqTusA as purified (-) and reduced with 5 mM DTT (+).

Recombinant HdTusA, TkTusA, TsTusA and AqTusA were purified in the absence of reducing agents. Mass spectrometry verified the masses without initiator methionine for all recombinant TusA proteins (Tables 1, S2). AqTusA was found to be glutathionylated to a small extent. This is likely a heterologous production artifact because the genes for the enzymes for biosynthesis of glutathione do not occur in the *Aq. aeolicus* genome.^(37)^ Upon native PAGE under non-reducing conditions, at least two bands were observed for all TusAs. In accordance, McTusA has been reported to occur both as monomers and dimers.^(15)^ Reduction with DTT resulted in the formation of faster migrating bands partially for AqTusA and TkTusA and completely for HdTusA and TsTusA. These bands likely represent monomeric TusA.

To elucidate the capacity of TusA from our four sHdr-containing model sulfur oxidizers for sulfur mobilization from inorganic and organic sulfur compounds, the purified proteins were incubated with 5 mM polysulfide (^-^SS_n_S^-^), thiosulfate (S_2_O_3_^2-^), tetrathionate (S_4_O_6_^2-^), and oxidized glutathione (GSSG) and analyzed by MALDI-ToF mass spectrometry. All reacted with polysulfide, resulting in mass increases of 32 Da, 64 Da or 96 Da, corresponding to the addition of one to three sulfur atoms (Table 1, Table S2). After incubation with tetrathionate, mass increases corresponding to addition of one or two sulfur atoms, a thiosulfonate group (-SSO_3_^-^), or -SSSO_3_^-^ were detected. With oxidized glutathione, mass increases of 305 Da were observed for TkTusA, AqTusA and HdTusA, which corresponds to covalently bound glutathione. The mobilization abilities for the bacterial TusAs AqTusA, TkTusA, TsTusA and HdTusA differed from those reported for archaeal TusA, which only mobilized thiosulfonate from tetrathionate but not sulfane sulfur from polysulfide. Like McTusA,^(15)^ none of the studied bacterial TusA proteins were modified by thiosulfate.

**TABLE 1.** Sulfur loading of native and variant TusA and DsrE3 proteins. Numbers in parentheses represent mass increases. –, no modification. Further information is available in Table S2.

| Protein | Expected masses [Da] | Detected masses [Da] | Modification |
| --- | --- | --- | --- |
| HdTusA + Polysulfide | 9149 | 9149 | - |
| | | 9180 ($\Delta 31$) | -S |
| | | 9208 ($\Delta 60$) | -S<sub>2</sub> |
| | | 9238 ($\Delta 90$) | -S<sub>3</sub> |
| HdTusA Cys<sup>13</sup>Ser + Polysulfide | 9135 | 9135 | - |
| HdTusA + S<sub>4</sub>O<sub>6</sub><sup>2-</sup> | 9149 | 9150 | - |
| | | 9148 ($\Delta 34$) | -S |
| | | 9212 ($\Delta 62$) | -S<sub>2</sub> |
| | | 9262 ($\Delta 112$) | -S-SO<sub>3</sub> |
| | | 9296 ($\Delta 146$) | -S<sub>2</sub>-SO<sub>3</sub> |
| AqTusA + Polysulfide | 9610 | 9612 | - |
| | | 9643 ($\Delta 31$) | -S |
| AqTusA Cys<sup>17</sup>Ser + Polysulfide | 9595 | 9595 | - |
| AqTusA Cys<sup>54</sup>Ser + Polysulfide | 9595 | 9596 | - |
| | | 9627 ($\Delta 31$) | -S |
| AqTusA + S<sub>4</sub>O<sub>6</sub><sup>2-</sup> | 9610 | 9610 | - |
| | | 9642 ($\Delta 32$) | -S |
| | | 9724 ($\Delta 114$) | -S-SO<sub>3</sub> |
| | | 9755 ($\Delta 145$) | -S<sub>2</sub>-SO<sub>3</sub> |
| AqTusA Cys<sup>17</sup>Ser + S<sub>4</sub>O<sub>6</sub><sup>2-</sup> | 9595 | 9595 | - |
| AqTusA Cys<sup>54</sup>Ser + S<sub>4</sub>O<sub>6</sub><sup>2-</sup> | 9595 | 9597 | - |
| | | 9630 ($\Delta 33$) | -S |
| | | 9710 ($\Delta 113$) | -S-SO<sub>3</sub> |
| HdDsrE3C + Polysulfide | 15520 | 15524 | - |
| | | 15561 ($\Delta 38$) | -S |
| | | 15588 ($\Delta 64$) | -S<sub>2</sub> |
| | | 15621 ($\Delta 98$) | -S<sub>3</sub> |
| HdDsrE3C Cys<sup>83</sup>Ser + Polysulfide | 15504 | 15503 | - |
| | | 15535 ($\Delta 32$) | -S |
| HdDsrE3C Cys<sup>84</sup>Ser + Polysulfide | 15504 | 15503 | - |
| HdDsrE3C + S<sub>4</sub>O<sub>6</sub><sup>2-</sup> | 15520 | 15517 | - |
| | | 15610 ($\Delta 93$) | -S<sub>3</sub> |
| | | 15629 ($\Delta 112$) | -S-SO<sub>3</sub><sup>-</sup> |
| | | 15658 ($\Delta 141$) | -S<sub>2</sub>-SO<sub>3</sub><sup>-</sup> |
| HdDsrE3C + GSSG | 15520 | 15518 | - |
| | | 15824 ($\Delta 305$) | -SG |
| TkDsrE3B + Polysulfide | 17201 | 17205 | - |
| | | 17236 ($\Delta 31$) | -S |
| | | 17320 ($\Delta 15$) | -S-SO<sub>3</sub><sup>-</sup> |
| TsDsrE3B + Polysulfide | 16619 | 16619 | - |
| | | 16653 ($\Delta 34$) | -S |
| | | 16683 ($\Delta 64$) | -S<sub>2</sub> |
| | | 16736 ($\Delta 117$) | -S<sub>2</sub>-SO<sub>3</sub><sup>-</sup> |

To identify the sulfur-binding cysteine with certainty, the cysteine of the CPXP motif was replaced by serine in both, HdTusA and AqTusA, as was the C-terminal partially conserved cysteine of AqTusA. No additional peaks were observed when AqTusA Cys^17^Ser or HdTusA-Cys^13^Ser were incubated with sulfur compounds. AqTusA Cys^54^Ser reacted with polysulfide and tetrathionate just as wild type AqTusA. The cysteine of the CPXP motif was thus confirmed as the sulfur-binding cysteine.

### 2.4 Properties of sHdr-associated Rhd442

The ability of rDsr associated rhodanese Rhd_2599 to transfer sulfur to the TusA protein encoded next to its gene has already been demonstrated for *Ac. vinosum*^(8)^ and there is no reason to doubt that closely related enzymes from other sulfur oxidizers (Figure 3b) exert the same function. All these proteins are single domain rhodaneses featuring a classical CRXGC[R/T] motif.^(33)^ On the other hand, information about the putative single domain rhodanese Rhd442 encoded in the vicinity of hyphomicrobial type I *shdr* clusters (Figure 1) is limited. Rhd442 has been described as a domain fused to sulfide:quinone oxidoreductase (SQR), e.g. in *Cupriavidus pinatubonensis* and that domain (CpRhd442) was shown to have rhodanese activity.^(38)^ The activity depended on the cysteines residue in the CRXGXR motif that the domain shares with classical rhodanese.^(33,39)^ A sequence reminiscent of that motif is also present in sHdr-associated Rhd442 (Figure 5a). The Alphafold models of HdRhd442 and CpRhd442 are similar (Figure 5b) and there is also high consistency with the crystal structure of a non-classical phosphatase from *Neisseria meningitidis* (PDB 2F46^(40)^) (Figure 5c). Recombinant HdRhd442 catalyzed sulfur transfer from thiosulfate to cyanide with a maximum specific activity of 360 mU/mg in the assay described by Ray et al. ^(41)^ Furthermore, the protein proved able to mobilize sulfur from polysulfide as shown by characteristic mass increases (Figure 5 d,e).

**FIGURE 5.**
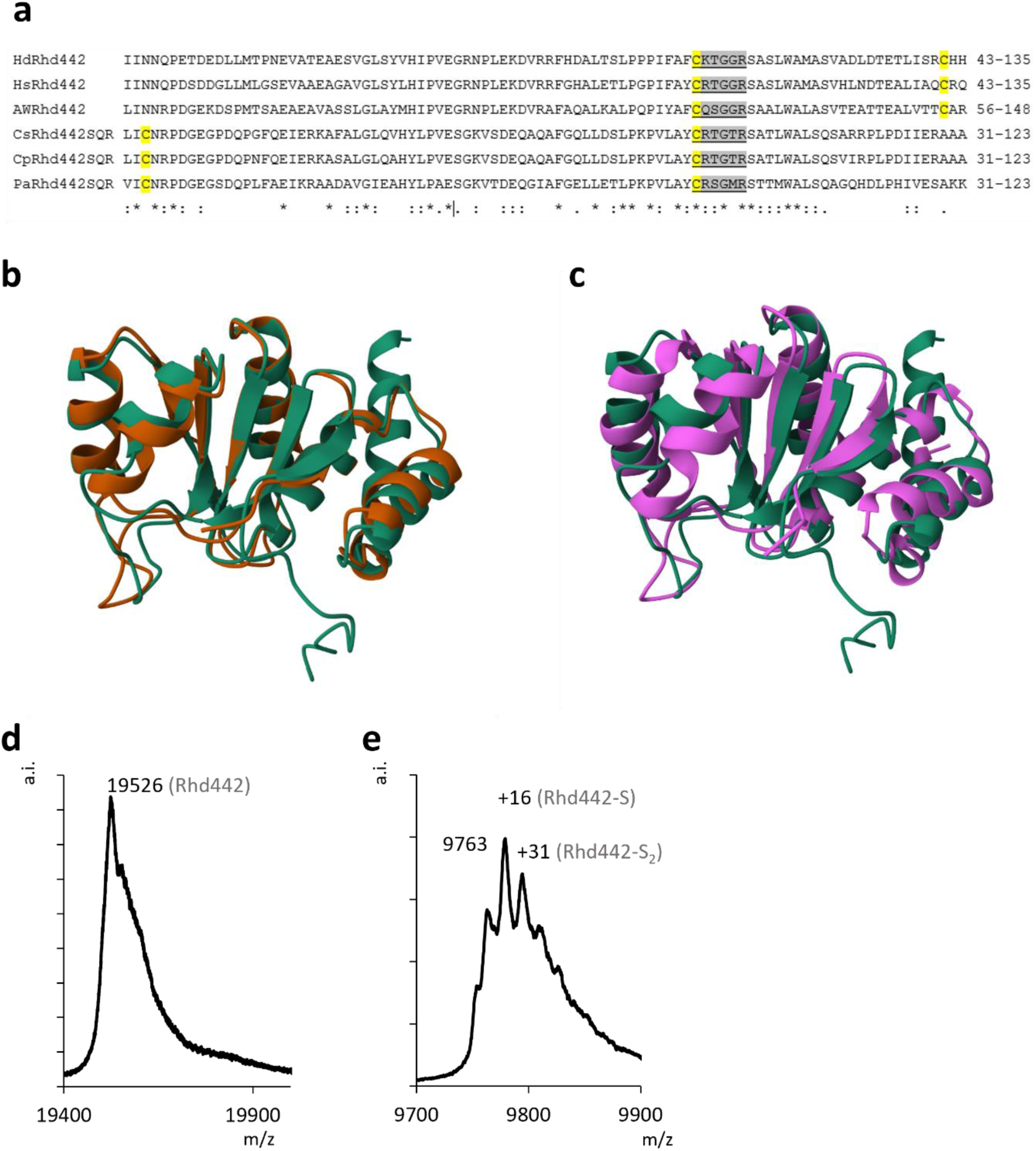
(a) Alignment of single domain Rhd442 from *Hm. denitrificans* (Hd), *Hyphomicrobium* sp018242215 (Hs), and AWTP1-13-sp008933705 (AW) with Rhd442 domains fused to sulfide:quinone oxidoreductase (SQR) from *Cupriavidus* sp. amp6 (Cs), *Cupriavidus pinatubonensis* (Cp), *Pseudomonas veronii* (Pa).^(38)^ Partially and fully conserved cysteines are highlighted in yellow. Sequences conforming to the rhodanese active site sequence logos^(39)^ are shaded in gray. Asterisk, fully conserved residues; colon, conservation between groups of strongly similar properties; dot, conservation between groups of weakly similar properties. (b) HdRhd442 Alphafold structure (green) overlayed with Rhd442 domain of *Cv. pinatubonensis* SQR (brown) and (c) non-classical phosphatase from *Ns. meningitidis* (PDB 2F46, purple). (d) Mass spectrum of recombinant HdRhd442 as isolated. (e) HdRhd442 mass spectrum after incubation with 0.5 mM polysulfide. Note that a distinguishable signal for Rhd442 and its persulfurated species was only observable for the double ionized protein.

### 2.5 Function and properties of sHdr-associated DsrE proteins

Analysis of recombinant DsrE3B from *Thioalkalivibrio sp.* K90mix and DsrE3C from *Hm. denitrificans* by gel permeation chromatography showed two peaks, one corresponding to the monomeric size and the other corresponding to a trimer (Figure 6a). Applying HdDsrE3C to native PAGE revealed a ladder of bands corresponding to higher oligomers (Figure 6b). Reduction of DsrE3C with 5 mM DTT resulted in a shift of the band pattern towards lower oligomers in relation to the non-reduced protein (Figure 6b). The structure of the trimeric complex was modelled by Alphafold (Figure 6c). Notably, the attempt to predict a hexamer for HdDsrE3C resulted in a complex which oligomerized by protein-protein interaction at the surface of only two subunits of each trimer, leaving two subunits for further docking with other units.

**FIGURE 6.**
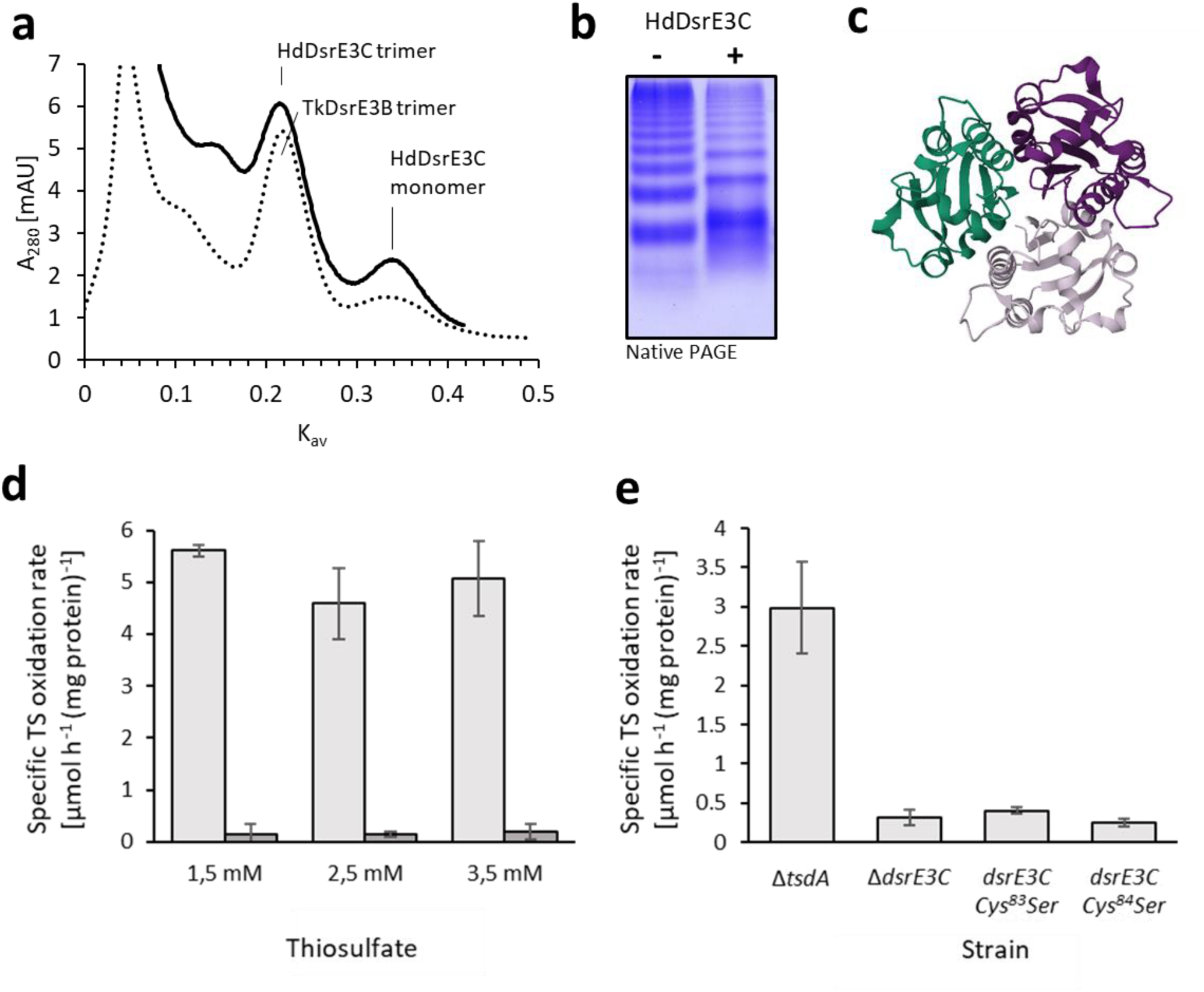
Function and properties of DsrE3B and DsrE3C proteins. (a) Gel permeation chromatography of HdDsrE3C and TkDsrE3B on Hiload 16/60 Superdex 75. (b) Native-PAGE of 3.5 µg HdDsrE3C as purified (-) and reduced with 5 mM DTT (+). (c) Alphafold model of the HdDsrE3C trimer. (d) Specific thiosulfate (TS) oxidation rates for *Hm. denitrificans* Δ*tsdA* (light gray columns) and *Hm. denitrificans* Δ*tsdA* Δ*dsrE3C* (dark gray columns) at the indicated initial thiosulfate concentrations. The corresponding growth curves are provided in Figure S1. Precultures contained 2 mM thiosulfate. (e) Specific thiosulfate (TS) oxidation rates of *Hm. denitrificans* Δ*tsdA* compared to a strain lacking the complete *dsrE3C* gene and two strains carrying *dsrE3C* genes encoding the indicating cysteine to serine exchanges grown with 2.5 mM thiosulfate. Note that specific thiosulfate oxidation rates are not fully comparable to the experiments shown in (d) because the growth experiments shown here were performed in a plate reader.

Recombinant DsrE3C and DsrE3B had masses corresponding to the polypeptides without the N-terminal starting methionine. In all cases of DsrE3B, modified species were detected that correspond to glycosylated derivatives (+178 Da) (Table 1, Figure S3, Figure 7b). This is due to the well documented addition of glucose to the His-tag of the recombinant proteins produced in *E. coli*.^(42)^ Sulfur mobilization assays showed reaction of HdDsrE3C with polysulfide, tetrathionate, and GSSG but not with thiosulfate. The DsrE3B proteins were persulfurated upon incubation with polysulfide. Oxidized species also occurred, probably due to the presence of oxygen. DsrE3C from *Hm. denitrificans* has an additional cysteine (Cys^83^) residing right next to the conserved cysteine (Cys^84^). To unambiguously identify the active site sulfur binding cysteine, both cysteines of HdDsrE3C were replaced with serine. While the Cys^83^Ser mutation did not significantly affect the sulfur binding properties of HdDsrE3C, the HdDsrE3C Cys^84^Ser variant was no longer persulfurated by polysulfide. In conclusion, Cys^84^ of HdDsrE3C was identified as the sulfur-binding cysteine.

**FIGURE 7.**
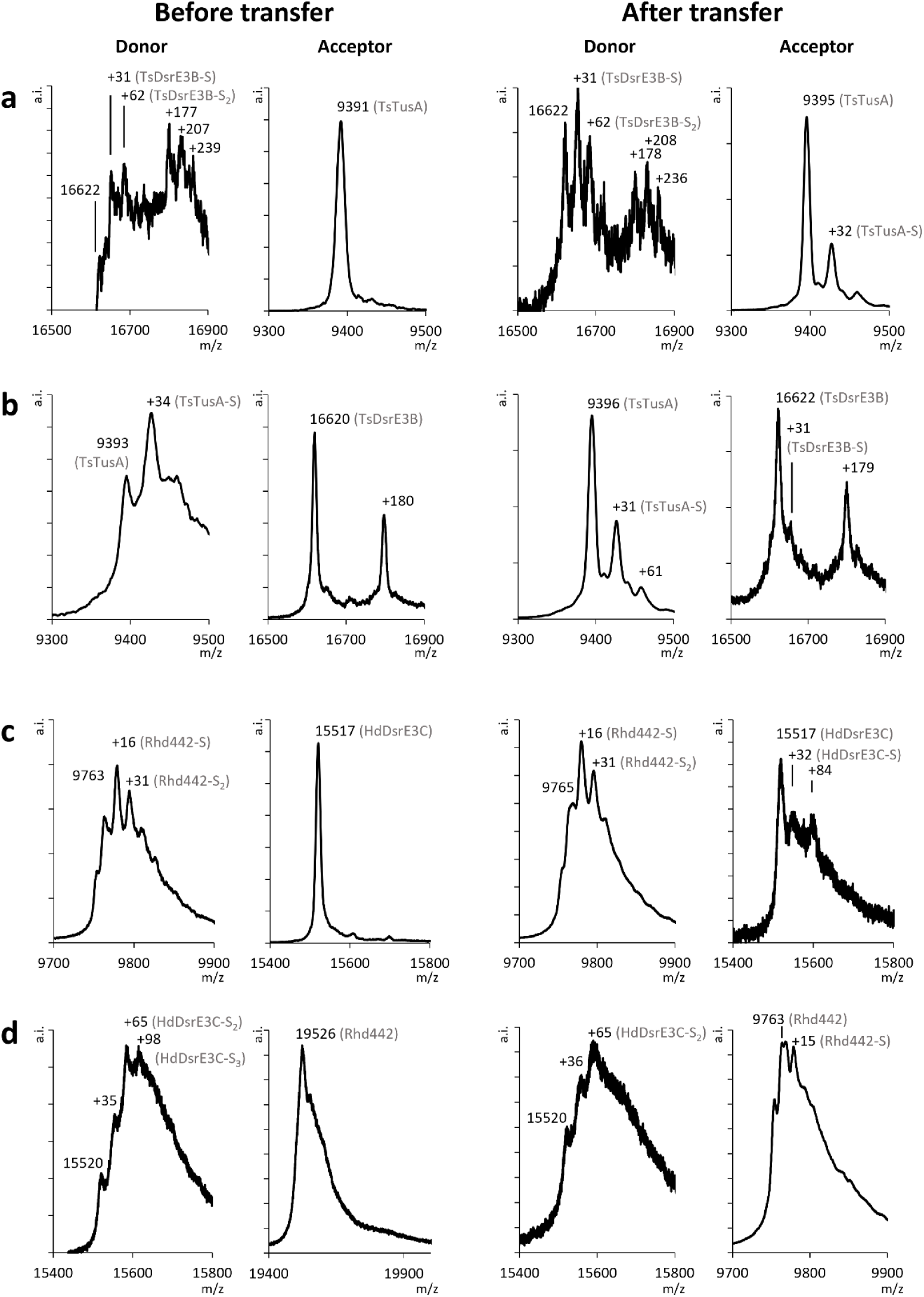
Sulfur transfer between TusA, DsrE3 proteins and Rhd442. Sulfur transfer reaction between TusA and DsrE3B from *Ts. sibirica* and between Rhd442 and DsrE3C from *Hm. denitrificans* are shown as examples. The full set of experiments is available as Figures S2, S3 and S4. (a) Left panels: TsDsrE3B as persulfurated donor after treatment with polysulfide and unmodified reduced TsTusA as acceptor; right panels: DsrE3B (donor) and TusA (acceptor) after the transfer reaction. The TsDsrE3B species exhibiting additional 178 Da are due to glucosylation of the His-Tagged protein.^(42)^ (b) Left panels: TsTusA as persulfurated donor after treatment with polysulfide and unmodified reduced TsDsrE3B as acceptor; right panels: TsTusA and TsDsrE3B after the transfer reaction. (c) Left panels: HdRhd442 as persulfurated donor after treatment with polysulfide and unmodified reduced HdDsrE3C as acceptor. Right panels: HdRhd442 (donor) and HdDsrE3C (acceptor) after the transfer reaction. (d) Left panels: HdDsrE3C as persulfurated donor after treatment with polysulfide and unmodified reduced HdRhd442 as acceptor. Right panels: HdDsrE3C (donor) and HdRhd442 (acceptor) after the transfer reaction.

*Hm. denitrificans* is accessible to genetic manipulation and indeed, the analysis of a mutant strain lacking *dsrE3C* provided important information. The deletion was established in the *Hm. denitrificans* Δ*tsdA* reference strain^(12,43)^ that completely oxidizes thiosulfate via the sHdr-pathway (Figure 6d, Figure S1). Compared with the reference strain, *Hm. Denitrificans* Δ*tsdA* Δ*dsrE3C* oxidized thiosulfate with a significantly decreased specific oxidation rate (Figure 6d). Thus, DsrE3C is crucial for the functionality of the sHdr system. In addition, the importance of the conserved Cys^84^ and the non-conserved Cys^83^ of HdDsrE3C was studied *in vivo* by replacing them with serine. Both exchanges resulted in significantly decreased specific thiosulfate oxidation rates compared to the reference strain (Figure 6e). We conclude that, although it is not involved in sulfur binding, Cys^83^ is important for DsrE3C function *in vivo*.

### 2.6 Interactions of sHdr-associated sulfurtransferases

The persulfurated DsrE3 proteins from our proteobacterial model organisms *Ts. sibirica*, *Thioalkalivibrio* sp. K90 and *Hm. denitrificans* were used as donors for the TusA proteins from the same organism, resulting in efficient sulfur transfer as shown by mass spectrometry (Figure 7a, Figure S2a, Figure S3a, Table S3). In contrast, in the opposite direction, when TusA acted as donor for the DsrE3 proteins, only a small amount of sulfur was added, if any was added at all (Figure 7b, Figure S2b, Figure S3b, Table S3). These *in vitro* results point at DsrE3 proteins transferring sulfur to TusA *in vivo,* whereas the opposite direction is unfavorable. Neither recombinant HdTusA-Cys^13^Ser nor HdDsrE3C-Cys^84^Ser accepted sulfur from the native persulfurated donor proteins, whereas HdDsrE3C-Cys^83^Ser accepted sulfur from persulfurated HdTusA (Table S3). The Cys^83^Ser mutation did not significantly affect the sulfur transfer properties of HdDsrE3C, while the HdDsrE3C Cys^84^Ser variant was neither persulfurated by polysulfide (see above) nor by HdTusA.

Persulfurized HdRhd442 was tested as sulfur donor for HdTusA and HdDsrE3C. Sulfane sulfur from HdDsrE3C was efficiently transferred to HdRhd442 (Figure 7c). Transfer from HdDsrE3C to HdRhd442 was also efficient (Figure 7d). With HdTusA as the acceptor molecule for HdRhd442, the transfer was inefficient as the intensity of the signal corresponding to the sulfurated species was low compared to the signal for the unmodified protein (Figure S4a). Transfer in the opposite direction was much more efficient (Figure S4b).

### 2.7 TusA and DsrE3C, DsrE3B and DsrE2 from *Aq. aeolicus* interact

*Aq. aeolicus* is an excellent model organism for identifying interactions between sHdr-associated sulfur transferases and other proteins. Membrane fractions, cell extracts and partially purified proteins from this hyperthermophile have repeatedly served as the basis for cross-linking, co-purification and co-migration approaches, which have enabled the identification of physiological protein partners.^(16,34,44)^ Here, we incubated pure recombinant His-tagged AqTusA with *Aq. aeolicus* soluble extracts, prepared from cells that had been grown in the presence of various ratios of hydrogen and thiosulfate,^(16)^ at room temperature as specified in the Material and methods section. In each case, after re-purification, AqTusA showed a mass gain of 32 Da, indicating a bound sulfane sulfur. This may be attributed to a transfer of sulfur originating from the cell extract since the heterologously produced AqTusA exhibited no such masses after purification (Table S1). In the first experiment, proteins interacting with TusA after 10 min incubation were assessed through mass spectrometry after re-purification and native PAGE (Figure 8a). This approach confirmed co-purification of TusA with AqDsrE3C (aq_390) and AqLbpA (aq_402) alongside 12 additional proteins (Table S4). Other sulfur transferases were not identified. In the second trial, AqTusA and *Aq. aeolicus* cell extract were incubated at room temperature for thirty minutes, followed by the identification by mass spectrometry of all proteins recovered after affinity chromatography. Cell extract without the addition of His-tagged TusA served as negative control. 253 proteins were found in the sample but not in the negative control and were therefore considered to be co-purified with AqTusA. Among them, AqDsrE3C (aq_390) was identified with one of the highest scores (Table S5). AqDsrE2A (aq_389) also eluted specifically in the presence of TusA but was identified with lower score and peptide number. Note that some subunits of the sHdr complex were also captured in this experiment (and not in the control, Table S5), some of which were identified with very high scores (the HdrB1 and HdrB2 subunits). All the other sHdr subunits were identified but also present (albeit with lower scores and peptide numbers) in the control sample, together with LbpA2 (aq_402) (Table S5). To confirm the interaction of TusA with DsrE-type and LbpA proteins, samples were applied to a SDS PAGE after re-purification (Figure 8b). AqDsrE3C and AqLbpA2 were clearly detected by mass spectrometry in one band that contains also AqDsrE3B (Aq_401) and 25 additional proteins (band 3, lane 3, Figure 8 and Table S6). In another band, AqDsrE2A (aq_389) was found together with LbpA3 (aq_1657), a putative thiol peroxidase (aq_488), and 14 additional proteins (band 2, lane 3, Figure 8 and Table S6).

**FIGURE 8.**
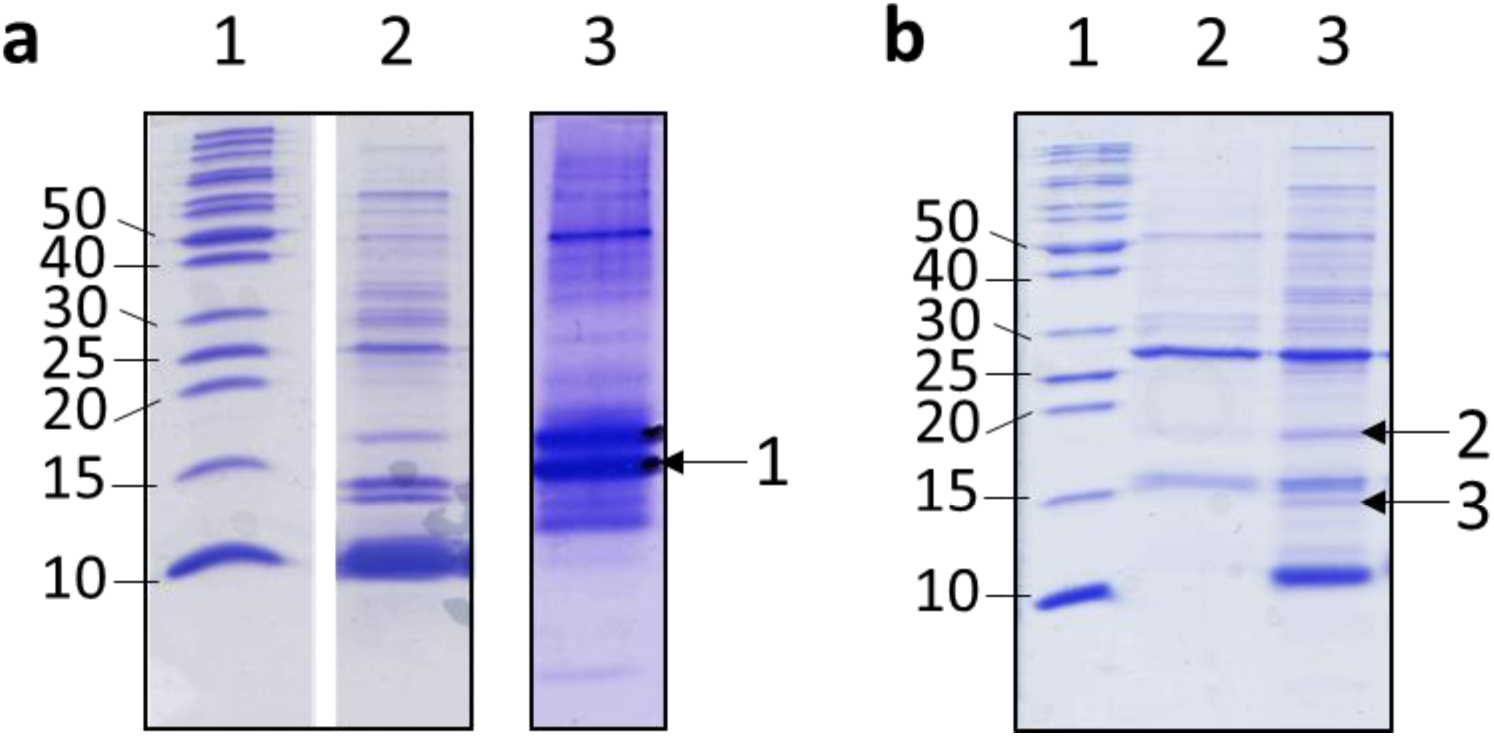
Re-purification experiments of AqTusA after incubated with *Aq. aeolicus* soluble extract in different conditions. (a) Proteins, eluted from the first re-purification column, were separated on a 18% SDS PAGE (lane 2) and a 15% native PAGE (lane 3). Six µg of total proteins were loaded in lanes 2 and 3. Proteins present in the band 1, indicated by an arrow, were identified by mass spectrometry (Table S4x). (b) Proteins, eluted from the second re-purification column, were separated on a 18% SDS PAGE. Lane 2 corresponds to the proteins retained in the control experiment (extract only) and lane 3 to the proteins potentially interacting with AqTusA. Same volumes of elution fractions were loaded in lanes 2 and 3 corresponding to 2 and 10 µg of total proteins, respectively. Proteins occurring in bands 2 and 3 were identified by mass spectrometry (Table S6). Molecular weight markers (kDa) for the SDS gels are shown in lanes 1.

## 3 DISCUSSION

When bound to a large molecule like a protein, persulfidic sulfane sulfur can be handled with high specificity.^(4)^ The sulfur transferases examined in this work perfectly illustrate this concept. Several independent lines of evidence have been combined to strongly suggest an important function for TusA- and DsrE3-type sulfur transferases in sulfur-oxidizing prokaryotes utilizing the sHdr pathway. First, the respective genes not only frequently occur in this physiological group (Figure 2), but in the majority of cases they co-localize with *shdr* genes (Figures 1 and 2), similar to what has been observed for the second well-established cytoplasmic oxidation pathway in sulfur-dependent lithotrophs, rDsr.^(8)^ Rhodanese Rhd442 is present in only a small number of sulfur oxidizers and therefore appears to be of minor importance. Second, deletion of the gene for DsrE3C from *Hm. denitrificans* resulted in a thiosulfate oxidation-negative phenotype, emphasizing the importance of the sulfur transferase. Third, the studied proteins all demonstrated the ability to bind sulfur upon incubation with inorganic polysulfide and transfer it to interaction partners, therefore acting as components of a sulfur trafficking cascade or network.

The investigated DsrE3B and DsrE3C proteins from bacteria were characterized as stable homotrimers in solution that have a tendency to assemble into higher oligomers, particularly hexamers. This is consistent with the available data on DsrE3A proteins from archaea. The homotrimeric form of the protein is observed in solution for the *Ms. cuprina* protein, and the *Saccharolobus solfataricus* (SSO1125) protein crystallizes as a trimer (PDB ID 3MC3). The DsrE2B sulfur transferases from the same archaea also form homotrimers (PDB ID 2QS7).^(15)^ In sulfur oxidizers utilizing the rDsr pathway, the heterohexameric DsrE_2_F_2_H_2_ complex plays a vital role in the sulfur relay system, which supplies the oxidizing enzyme rDsrAB. The subunits are arranged in two stacked DsrEFH rings, each resembling the trimeric rings observed in DsrE3A crystals (PDB ID 3MC3) and DsrE3C Alphafold models (Figure 6c). DsrE, DsrF and DsrH are related to each other.^(10)^ In summary, the formation of trimers is a general and common property of DsrE and DsrE3-type proteins, that has been conserved throughout evolution. It is probable, that the three distinct subunits in DsrEFH have evolved due to gene duplications and specialization of ancestral DsrE-type proteins. In fact, from an evolutionary perspective, the DsrE3 proteins are older than the DsrE in DsrEFH.^(48)^ Filamentation has not been reported for the DsrEFH complex nor for any other DsrE-type; it is a novel characteristic of DsrE3C from *Hm. denitrificans*. As previously stated, all DsrE-like proteins possess a single conserved cysteine residue (Cys^84^ in HdDsrE3C). The importance of the equivalent cysteine for sulfur binding and transfer has so far only been shown *in vitro* for the *Ms. cuprina* protein DsrE3A and both *in vivo* and *in vitro* for the distantly related DsrE from the *A. vinosum* DsrEFH complex. Here, we prove the crucial role of this residue through *in vivo* and *in vitro* studies on DsrE3C from *Hm. denitrificans.* Furthermore, we show that the adjacent Cys^83^ is important for DsrE3C function *in vivo*, even though it does not participate in sulfur binding.

Just like archaeal McDsrE3A,^(15)^ all DsrE3B, DsrE3C and TusA proteins from bacteria that were examined in this study displayed reaction with tetrathionate *in vitro*. However, it is impossible that thiosulfonates derived from tetrathionate are the physiologically relevant form of sulfur processed in the cytoplasm of these bacteria. Neither *Hm. denitrificans*^(12)^ nor *Thioalkalivibrio sp.* K90mix^(45)^, *Aq. aeolicus*^(49)^ or *Ts. sibirica*^(46)^ can metabolize tetrathionate. If we seek a type of sulfur that every organism studied can use as a cytoplasmic sulfur currency, then only polysulfide or sulfane sulfur is left, as *Ts. sibirica* cannot oxidize thiosulfate either (Figure 9). In addition, HdTusA and HdDsrE3C are unable to mobilize sulfane sulfur from thiosulfate *in vitro*.

**FIGURE 9.**
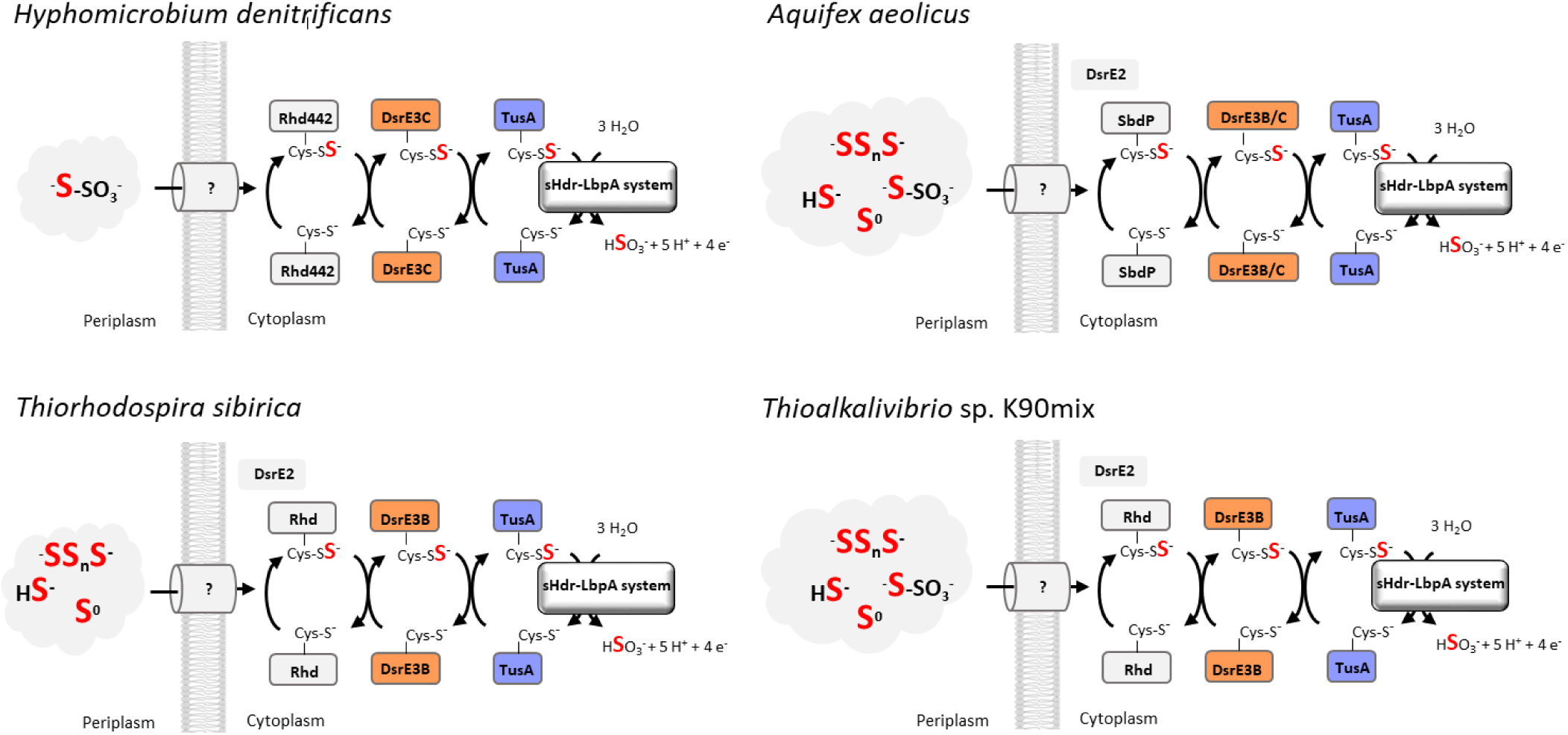
Sulfur relay systems in the four bacterial model organisms studied in this work. *Hm. denitrificans* oxidizes thiosulfate and dimethyl sulfide. Thiosulfate is an intermediate of DMS oxidation.^(11,12)^ *Aq. aeolicus*^(37)^ and *Thioalkalivibrio* sp. K90mix^(45)^ metabolize a wide range of inorganic sulfur substrates including thiosulfate, whereas *Ts. sibirica* is unable to oxidize thiosulfate.^(46)^ It was originally predicted that DsrE2A from *Aq. aeolicus* (Aq_389) is bound to the membrane via two transmembrane segments. However, recent deep-learning based programs like DeepTMHMM^(47)^ challenge this view not only for the *Aquifex* protein but also for the corresponding enzymes found in *Ts. sibirica* and *Thioalkalivibrio* sp. K90mix.

In dissimilatory sulfur oxidizers, the oxidation of sulfide and thiosulfate is always initiated outside of the cytoplasm.^(2,6)^ In many cases, sulfane sulfur is then imported into the cytoplasm where it is further oxidized (Figure 9). It is not yet clear how this import is accomplished. *Hm. denitrificans* possesses two candidate transporters (SoxT1A and SoxT1B, Figure 1) encoded in close proximity to the genes encoding Sox proteins that are involved in the initial steps of thiosulfate oxidation and the genes for the cytoplasmic sHdr system.^(43,50)^ However, evidence supporting the proposed sulfur transport has yet to be presented. Related transporters are not encoded in *Aq. aeolicus* and *Ts. sibirica* such that they cannot be of general importance. Rhodanese-like sulfurtransferases including Rhd442, with its comparatively narrow distribution, are potential primary sulfur acceptors and distributors in the cytoplasm. They occur in all examined organisms (Figures 1, 2, 9) and have well established sulfur transfer activity. HdRhd442 was shown here to interact with HdDsrE3C. This protein, in turn, was established as an indispensable component of the sulfur-handling cascade that feeds the type I sHdr system of *Hm. denitrificans. In vitro*, both HdDsrE3C and the DsrE3B proteins from gammaproteobacterial model organisms shuttle sulfane sulfur to TusA (Figures 7, 9). Efficient sulfur transfer to TusA from the same organism was observed with HdDsrE3C, TkDsrE3B and TsDsrE3B, while transfer in the opposite direction was either undetectable or comparatively inefficient. Unidirectional transfer has also been observed for DsrE3A and TusA from *Ms. cuprina*, where a thiosulfonate group is moved from DsrE3A to TusA but not vice versa.^(15)^ In the absence of a DsrE3C homolog, as in *Thioalkalivibrio* sp. K90mix, the sulfurtransferase DsrE3B may functionally substitute for DsrE3C. It is very important to note that each sulfur transferase may interact with multiple partners as illustrated by TusA from *Aq. aeolicus* which was purified along with three DsrE, two AqLbpA and several sHdr proteins. This finding is particularly significant as it establishes a direct link between sulfur transferases and the sHdr-LbpA complex where sulfane sulfur is oxidized to sulfite.^(11,12,18)^

## 4 CONCLUSIONS

We conclude that cytoplasmic sHdr systems for sulfane sulfur oxidation are always accompanied by sets of sulfur transferases. The exact composition of these sets may vary (Figure 9). *In vivo,* a strict unidirectional transfer of sulfur between the components is unlikely. Rather, it can be assumed that a network of sulfur-binding proteins exists, each with a pool of bound sulfur. Sulfur flux can be shifted in one direction or the other depending on the metabolic requirements. A single pair of sulfur-binding proteins with a preferred transfer direction, such as a DsrE3-type protein toward TusA, may well be sufficient to push sulfur into the sink where it is further metabolized or needed. Multiple possible interactions are most easily exemplified for the TusA protein. In organisms such as *Hm. denitrificans*, which differ from *E. coli* by containing just one *tusA* gene, the same protein must likely fulfill a variety of functions, including providing substrate for sulfur oxidation, and supplying sulfur for tRNA thiolation and biosynthesis of cofactors and Fe/S clusters.

## 5 MATERIALS AND METHODS

### 5.1 Bacterial strains, plasmids, primers, and growth conditions

Table S7 lists the bacterial strains, and plasmids that were used for this study. *Escherichia coli* strains were grown on complex lysogeny broth (LB) medium^(51)^ under aerobic conditions at 37°C unless otherwise indicated. *E. coli* 10β was used for molecular cloning. *E. coli* BL21 (DE3) was used for recombinant protein production. *Hm. denitrificans* strains were cultivated in minimal media kept at pH 7.2 with 24.4 mM methanol and 100 mM 3-(*N*-Morpholino) propanesulfonic acid (MOPS) buffer as described before.^(12,43)^ Thiosulfate was added as needed. Antibiotics for *E. coli* and *Hm. denitrificans* were used at the following concentrations (in μg ml^-1^): ampicillin, 100; kanamycin, 50; streptomycin, 200; chloramphenicol, 25.

### 5.2 Recombinant DNA techniques

Standard techniques for DNA manipulation and cloning were used unless otherwise indicated.^(52)^ Restriction enzymes, T4 ligase and Q5 polymerase were obtained from New England Biolabs (Ipswich, UK) and used according to the manusfacturer’s instructions. Oligonucleotides for cloning were obtained from Eurofins MWG (Ebersberg, Germany). Plasmid DNA from *E. coli* was purified using the GenJET Plasmid Miniprep kit (Thermo Scientific, Waltham, USA). Chromosomal DNA from *Hm. denitrificans, Ts. sibirica* and *Thioalkalivibrio* strains was prepared using the Simplex easy DNA kit (GEN-IAL GmbH, Troisdorf, Germany). Total DNA from 20 mg of *Aq. aeolicus* cells was extracted using the phenol-chloroform extraction method. Cells were resuspended in 150 µl of TEN buffer (10 mM Tris pH8, 150 mM NaCl, 10 mM EDTA) and 300 µl of SDS-EB (100 mM Tris pH8, 400 mM NaCl, 40 mM EDTA, 2% SDS). 2 µl of RNAse A solution (4 mg ml^-1^) (Promega) were added before incubation at 37°C for 15 min. 350 µl phenol/chloroform-isoamylalcohol (25:24:1 mixture, Biosolve) were added and the mixture was vortexed for 30 s and spun for 3 min at 14,000×g. The upper phase was transferred to a new tube with 300 µl of chloroform-isoamylalcohol, vortexed and spun again. The upper phase was then incubated with 600 µl isopropanol for 30 min at −25°C and spun at 4°C for 30 min. The DNA pellet was washed with 70% ethanol, left to air dry and dissolved in 100 µl H_2_O for 30 min at 65°C. 1 µl was used for 25 µl PCR reaction.

### 5.3 Construction of *Hm. denitrificans* mutant strains

For markerless deletion of the *Hm. denitrificans dsrE3C* (Hden_0688) gene by splicing overlap extension (SOE) (Horton, 1995), PCR fragments were constructed using the primers Hden0688_Up_Fw, Hden0688_Up_Rev, Hden0688_Down_Fw, Hden0688_Down_Rev (Table S1). The resulting 2.08 kb SOE PCR fragment was cloned into the XbaI and PstI sites of pK18mobscaB-Tc. The final construct pK18mobsacB_Tc_Δ*dsrE3C* was electroporated into *H. denitrificans* Δ*tsdA* and transformants were selected using previously published procedures ^(11,12)^. Single crossover recombinants were Cm^r^ and Tc^r^. Double crossover recombinants were Tc^s^ and survived in the presence of sucrose due to loss of both, the vector-encoded levansucrase (SacB) and the tetracycline resistance gene. For chromosomal integration of the genes encoding DsrE3C Cys^83^Ser and DsrE3C Cys^84^Ser, the modified genes and upstream as well as downstream sequences were amplified by SOE PCR using primers Hden0688_Up_Fw, Hden0688_Down_Rev, Hden0688_C83S_Fw, Hden0688_C83S_Rev, and Hden0688_Up_Fw, Hden0688_Down_Rev, Hden0688_C84S_Fw, Hden0688_C84S_Rev (Table S1), respectively. The final plasmids pk18*mobsacB-dsrE3C-C83S*-Tc and pk18*mobsacB-dsrE3C-C84S*-Tc were transferred into *Hm. denitrificans* Δ*tsdA* Δ*dsrE3C* and double crossover recombinants were selected as described previously.^(12)^ The genotypes of the *Hm. denitrificans* mutant strains generated in this study were confirmed by PCR.

### 5.4 Characterization of phenotypes and quantification of sulfur compounds

Growth experiments with *Hm. denitrificans* were run in in Erlenmeyer flasks with media containing 24.4 mM methanol and varying concentrations of thiosulfate as necessary.^(43)^ Thiosulfate and sulfite concentrations and biomass content were determined by previously described methods.^(43,53)^ All growth experiments were repeated three to five times. Representative experiments with two biological replicates for each strain are shown. All quantifications are based on at least three technical replicates. Alternatively, growth experiments were run in 48-well microtiter plates. Plates were continuously shaken at 200 rpm and growth was followed by measuring optical density at 600 nm every 5 min using an Infinite 200Pro (Tecan, Crailsheim, Germany) plate reader. Samples for thiosulfate determination were taken as previously described.^(43)^

### 5.5 Cloning, site-directed mutagenesis, overproduction, purification and size exclusion chromatography of recombinant proteins

The 378-bp *dsrE3C* gene was amplified from *Hm. denitrificans* genomic DNA with the primers Hden0688 (*dsrE3C*)_NdeI_fw and Hden0688 (*dsrE3C*)_BamHI_rev (Table S1) and cloned between the NdeI and BamHI sites of pET15b(+), resulting in pET15b-Hd-DsrE3C. Analogous procedures were followed for *dsrE3B* from *Thioalkalivibrio* sp. K90mix which was cloned between the NdeI and BamHI sites and *dsrE3B* from *Ts. sibirica* which was cloned between the XhoI and NdeI sites generating the plasmids pET15b-TkDsrE3B and pET15b-TsDsrE3B. The *tusA* genes were amplified from *Hm. denitrificans*, *Thioalkalivibrio* sp. K90mix or *Ts. sibirica* genomic DNA with primers adding a sequence for a C-terminal Strep-tag and cloned between the NdeI and EcoRI sites of pET-22b(+). The *tusA* gene from *Aq. aeolicus* (aq_388a, coding for the protein WP_024015099.1) was amplified from genomic DNA with primers Aq388a_NdeI_fw and Aq388a_XhoI_rev (Table S1) introducing at the C-terminal position of the protein the two amino acids Leu and Glu directly followed by a 6His-tag, and cloned into the pET24a expression plasmid to generate the plasmid pET24a-AqTusA. Cysteine to serine exchanges were implemented to HdDsrE3C by SOE PCR using primers sets Hden0688_NdeI_fw and Hden0688_BamHI_rev, Hden0688_C83S_Fw, Hden0688_C83S_Rev and Hden0688_NdeI_fw and Hden0688_BamHI_rev, Hden0688_C84S_Fw, Hden0688_C84S_Rev, respectively, resulting in plasmids pET15b-DsrE3C-C83S and pET15b-DsrE3C-C84S. A cysteine to serine exchange was introduced into *Hm. denitrificans* TusA using the same method and the primers listed in Table S7. The QuikChange site-directed mutagenesis Kit (Stratagene) was used to generate the *Aq. aeolicus tusA* mutated genes (coding for AqTusA Cys^17^Ser and AqTusA Cys^54^Ser) with the primers Aq388a_C17S_fw, Aq388a_C17S_rev and Aq388a_C54S_fw, Aq388a_C54S_rev using the pET24a-AqTusA plasmid.

Recombinant DsrE3B, DsrE3C and TusA proteins were produced in *E. coli* BL21(DE3). Overnight precultures were used to inoculate fresh LB medium with a ratio of 1:50 (v/v). Synthesis of recombinant proteins was induced by the addition of 0.1 or 1mM (for AqTusA) IPTG when cultures had reached an OD600 of 0.6–0.8, followed by incubation for 2.5 h at 37 °C. Cells were harvested by centrifugation (11,000 × g, 20 min, 4°C). Strep-tagged proteins were resuspended in 50 mM Tris-HCl buffer (pH 7.5) containing 150 mM NaCl. His-tagged proteins were resuspended in buffer containing 20 mM sodium-phosphate (20 mM Tris-HCl for AqTusA), 500 mM NaCl and 50 mM imidazole (pH 7.4). Cells were lysed by sonication (or with a cell disruptor for AqTusA). Insoluble cell material was subsequently removed by centrifugation (16,100 × g, 30 min, 4°C). His-tagged and Strep-tagged proteins were purified with Ni-NTA Agarose (Jena Bioscience) (except for AqTuA that was purified with a HiScreen Ni FF column (Cytiva)) and Strep-Tactin Superflow (IBA Lifesciences, Göttingen, Germany), respectively, according to the manufacturer’s instructions. The proteins were then transferred to salt-free 50 mM Tris-HCl buffer (pH 7.5) or to 20 mM Tris-HCl buffer pH 7.4 for AqTusA and stored at −70°C. Size exclusion chromatography on HiLoad 16/60 Superdex™ 75 (Cytiva, Freiburg, Germany) was performed as described in Li et al.^(50)^

### 5.6 Protein-protein interaction in cell extracts

To detect interaction between AqTusA and DsrE-like proteins, TusA from *Aq. aeolicus* (200 μg) was incubated with crude soluble extract of *Aq. aeolicus* at room temperature and re-purified via His tag affinity-chromatography (HisTrap FF 1 ml column, Cytiva) according to the manufacturer’s instructions. Soluble extracts were prepared as previously described^(49)^ except that cells were resuspended in 50 mM Tris-HCl pH 7.3. Two different experiments were run: in the first one, purified AqTusA was incubated for 10 min with 500 µl of an extract obtained from cells grown with excess of hydrogen in the presence of thiosulfate (called 100% H_2_) as described previously;^(16)^ in the second trial, the protein was incubated for 30 min with 2 ml of extract prepared from cells grown with a lower amount of H_2_ in the presence of thiosulfate (referred as 30% H_2_, condition in which the Hdr amount in cells is higher^(16)^). In both cases, the AqTusA-extract mixtures were immediately frozen in liquid nitrogen after incubation and thawed just prior to purification. Proteins were eluted with a buffer containing 50 mM Tris-HCl pH 7.3, 150 mM NaCl and 250 mM imidazole. A control experiment was run with the same extract but without TusA. After dialysis on a Vivaspin concentrator (molecular mass cutoff 3000 Da) with 50 mM Tris HCl pH 7.3, eluted proteins from both columns underwent 18% SDS PAGE or 15% Tris-Glycine native PAGE (without any reducing agent).^(54)^ Resulting bands were cut out of the gel and analyzed by LC-MSMS as described previously.^(16)^ For the second experiment, total proteins in the elution fraction of the column were identified after the proteins were introduced in a 5% acrylamide stacking gel, as described in Prioretti et al.^(44)^ under the name “stacking method”. Protein concentrations were determined with the BCA protein assay kit from Sigma-Aldrich.

### 5.7 Sulfur binding and transfer experiments

For sulfur binding experiments with the recombinant sulfur transferases from *Hm. denitrificans*, *Thioalkalivibrio* sp. K90mix and *Ts. sibirica*, 1.5 nmol of the proteins were incubated with 5 mM polysulfide, thiosulfate, tetrathionate or GSSG in 20 µl 50 mM Tris–HCl pH 7,5. The polysulfide stock solution needed for these experiments was prepared, diluted and used as previously described.^(50,55)^ For sulfur transfer experiments, 1.5 nmol of the putative sulfur-donating protein were incubated with 0.5 mM polysulfide for 30 minutes at room temperature in 20 µl 50 mM Tris–HCl pH 7,5. Acceptor proteins were reduced with 1 mM DTT under the same conditions. Excess of polysulfide or DTT was removed with micro Bio-Spin 6 columns (GE Healthcare, Munich, Germany) equilibrated with 50 mM Tris-HCl pH 7.5. Efficient removal of polysulfide was confirmed by adding a filtered protein-free polysulfide solution to an acceptor protein, which was then tested for persulfuration via mass spectrometry. Donor and acceptor proteins were mixed in a 1:1 ratio to a final volume of 40 µl and incubated for 30 min at room temperature. The mixtures were then stored at −70°C. For mass spectrometry, samples of 20 µl were desalted by ZipTipC4 Pipette tips (Merck Millipore, Darmstadt, Germany), crystallized in a 2’,6’-dihydroxyacetophenon matrix and measured by MALDI-ToF (matrix-assisted laser desorption ionization-time-of-flight) mass spectrometry at the Core Facility Protein synthesis & BioAnalytics, Pharmaceutical Institute, University of Bonn as previously described.^(54)^

For sulfur loading of AqTusA, 2 µg of protein was incubated for 30 min at 65°C, with 5 mM (final concentration) of the tested sulfur compound, in a final volume of 5 µl in 50 mM Tris-HCl pH 7.3. If reduction was desired, samples were incubated, with 10 mM DTT (final concentration) for 45 min at room temperature after the sulfur loading. Samples were immediately frozen in liquid nitrogen and stored at −80°C until further use. Before MALDI-ToF mass spectrometry analysis, the samples were desalted and concentrated with ZipTip C18 (Merck Millipore) using 0.1% trifluoroacetic acid (TFA) as desalting solution and 70% acetonitrile/0.1% TFA as elution solution. 1 μl of samples (∼41 pmol) mixed with 1 μl of matrix α-cyano-4-hydroxycinnamic acid were analyzed using the mass spectrometer Microflex II (Bruker). Three µg of AqTusA, incubated with cell extracts and re-purified on a HisTrap column (see paragraph 5.6), was diluted in 50 mM Tris-HCl pH 7.3 in a volume of 5 µl, desalted with ZipTipC18 and analyzed by MALDI-TOF.

### 5.8 Generation of datasets for phylogenetic and similarity network analyses

Archaeal and bacterial genomes were downloaded from Genome Taxonomy Database (GTDB, release R207). In GTDB, all genomes are sorted according to validly published taxonomies, they are pre-validated and have high quality (completeness minus 5*contamination must be higher than 50%). One representative of each of the current 65,703 species clusters was analyzed. Open reading frames were determined using Prodigal^(56)^ and subsequently annotated for sulfur related proteins via HMSS2.^(21)^ Annotation was extended by HMMs from TIGRFAMs^(57)^ and Pfam^(58)^ databases representing the 16 syntenic ribosomal proteins RpL2, 3, 4, 5, 6, 14, 15, 16, 18, 22, and 24, and RpS3, 8, 10, 17, and 19. A type I sHdr system was considered to be present if the core genes *shdrC1B1AHC2B2* were present in a syntenic gene cluster. For a type II sHdr system gene cluster *shdrC1B1AHB3* and *etfAB* had to be present in a single syntenic gene cluster.^(11,14)^

### 5.9 Phylogenetic tree inference and structural modelling

For phylogenetic tree inference, proteins were aligned using MAFFT^(59)^ and trimmed with BMGE^(60)^ (entropy threshold = 0.95, minimum length = 1, matrix = BLOSUM30). Alignments were then used for maximum likelihood phylogeny inference using IQ-TREE v1.6.12^(61)^ implemented on the “bonna” high performance clusters of the University of Bonn. The best-fitting model of sequence evolution was selected using ModelFinder.^(62)^ Branch support was then calculated by SH-aLRT (2000 replicates),^(63)^ aBayes (2000 replicates)^(64)^ and ultrafast bootstrap (2000 replicates).^(65)^ Finally, trees were displayed using iTol.^(66)^ For species tree inference, results for each ribosomal protein were individually aligned, trimmed and subsequently concatenated before they were used for phylogenetic tree construction. Structural models of proteins and protein complexes were generated using Alphafold2.^(67)^

### 5.10 Sequence similarity network analysis

Amino acid sequences for the sequence similarity network were derived from the search of the GTDB dataset with HMSS2.^(21)^ Groups were chosen based on their sequence similarity and genomic context, which depended on the specific question being investigated. From all selected sequences a meaningful and diverse set was derived via dereplication with mmseqs2 linclust^(68)^with default settings. Sequence similarity analysis was performed by an all versus all comparison with mmseqs2 search.^(68,69)^ The similarity matrix was modified in cytoscape and edged were filtered stepwise until optimal clustering was observed. This status was characterized as the minimum number of clusters while maintaining the highest number of connectivity within a group of annotated proteins, visualizing the grouping of proteins on a deep branching level.

## ACKNOWLEDGEMENTS

This work was funded by the Deutsche Forschungsgemeinschaft (Grant Da 351/13-1). Tomohisa Sebastian Tanabe received a scholarship from the Studienstiftung des Deutschen Volkes. We gratefully acknowledge the access to the Bonna HPC cluster hosted by the University of Bonn along with the support provided by its High Performance Computing & Analytics Lab. We gratefully thank for the support of France-Germany Hubert Curien Procope program for its exchange funding (n° 40444VM). We gratefully acknowledge the support of the Core Facility “Protein Synthesis and Bioanalytics” of the University of Bonn. The authors are also grateful to Régine Lebrun from the Core Facility “Protein Synthesis Mass spectrometry and Bioanalytics” proteomic platform of the University Mediterranean Institute of Microbiology (IMM) in Marseille (France), which is a part of the network “Marseille Protéomique”, IBiSA, and the Proteomic Platform IBiSA of the IMM, CNRS, Aix-Marseille Université for performing mass spectrometry. We thank Souhéla Boughanemi and Robert van Lis (BIP-CNRS, Marseille) for *Aq. aeolicus tusA* cloning and site directed mutagenesis. We thank Alina Ballas for help with *Hm. denitrificans* growth experiments.

## SUPPLEMENTARY MATERIAL

Additional supporting information may be found online in the supporting Information section at the end of this article

