## Supplementary material for "A cascade of sulfur transferases delivers sulfur to the sulfur-oxidizing heterodisulfide reductase-like complex": Figures S1-S4, Tables S1-S7: Figure S1 Growth experiments DsrE strains.docx

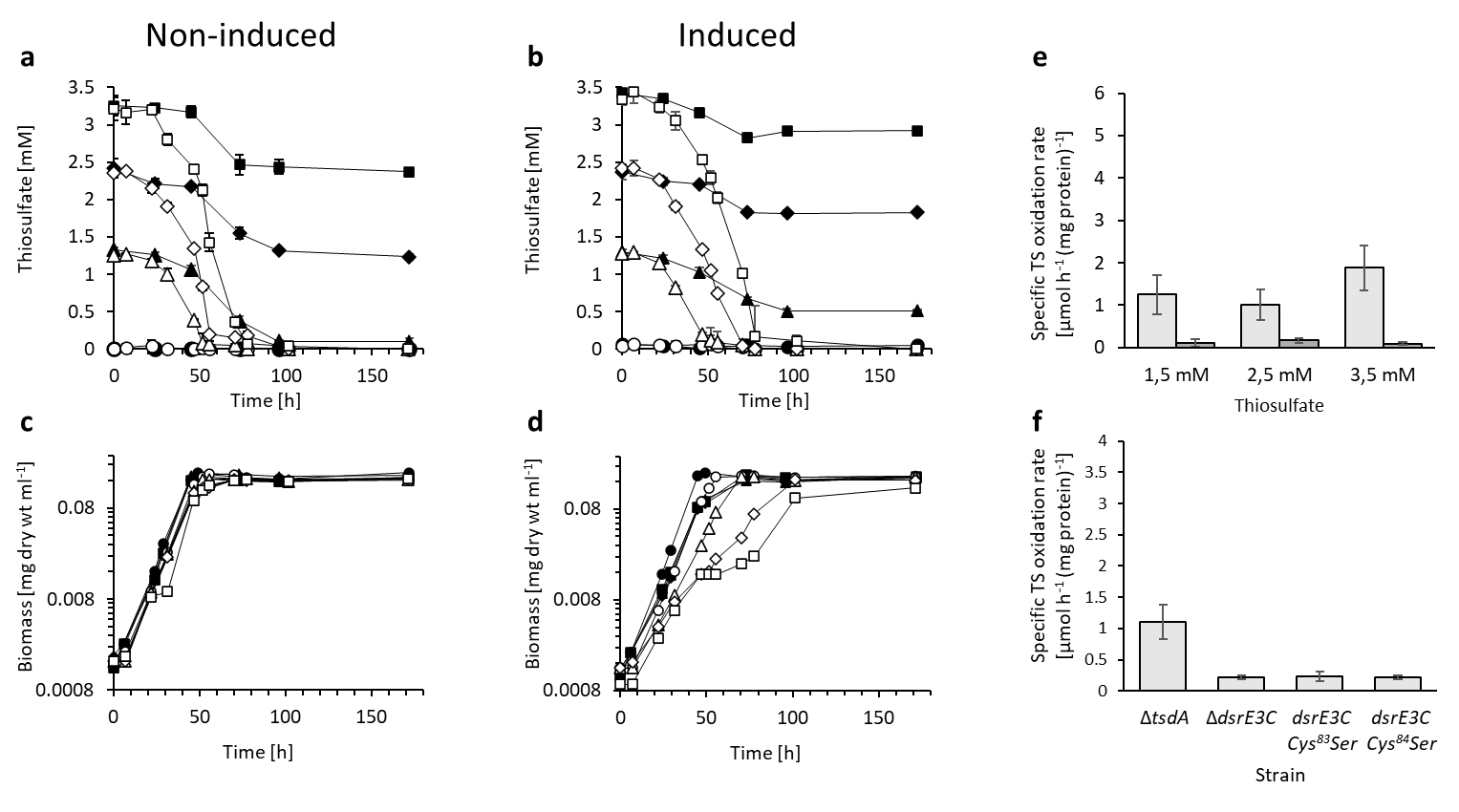

**FIGURE S1.** Thiosulfate consumption (a,b) and growth (c,d) band of *H. denitrificans* Δ*tsdA* (open symbols) and *H. denitrificans* Δ*tsdA* Δ*dsrE3* (filled symbols) in a medium with 24.4 mM methanol and different thiosulfate concentrations (3.5 mM, boxes, 2.5 mM, diamonds, 1.5 mM, triangles, no thiosulfate, circles). Precultures contained either no thiosulfate (non-induced, panels a and c) or 2 mM thiosulfate (induced, panels b and d). In general, induced strains oxidized thiosulfate at a higher specific oxidation rate than non-induced strains. When the Δ*tsdA* reference strain is grown with thiosulfate as an additional electron source, it excretes toxic sulfite, which causes growth retardation (J. Li, Koch, et al., 2023). Functionality of the sHdr-LbpA pathway is thus easily detectable. Growth inhibition of the reference strain was proportional to the initial thiosulfate concentration. Growth retardation was not observed for strain *H. denitrificans* Δ*tsdA* Δ*dsrE3C*. (e) Specific thiosulfate oxidation rate for the non-induced strains *H. denitrificans* Δ*tsdA* (light gray) and *H. denitrificans* Δ*tsdA* Δ*dsrE3C* (dark gray). (f) Specific thiosulfate oxidation rates of *H. denitrificans* Δ*tsdA* compared to the mutant lacking the *dsrE3C* gene and two strains carrying *dsrE3C* genes encoding the indicating cysteine to serine exchanges. Cultures were grown with 2.5 mM thiosulfate and without prior induction. Note that specific thiosulfate oxidation rates are not fully comparable to the experiments shown in (e) because the growth experiments shown here were performed in a plate reader.
