## Supplementary material for "A cascade of sulfur transferases delivers sulfur to the sulfur-oxidizing heterodisulfide reductase-like complex": Figures S1-S4, Tables S1-S7: Figure S2 Transfer_HdTusA_HdDsrE3C.docx

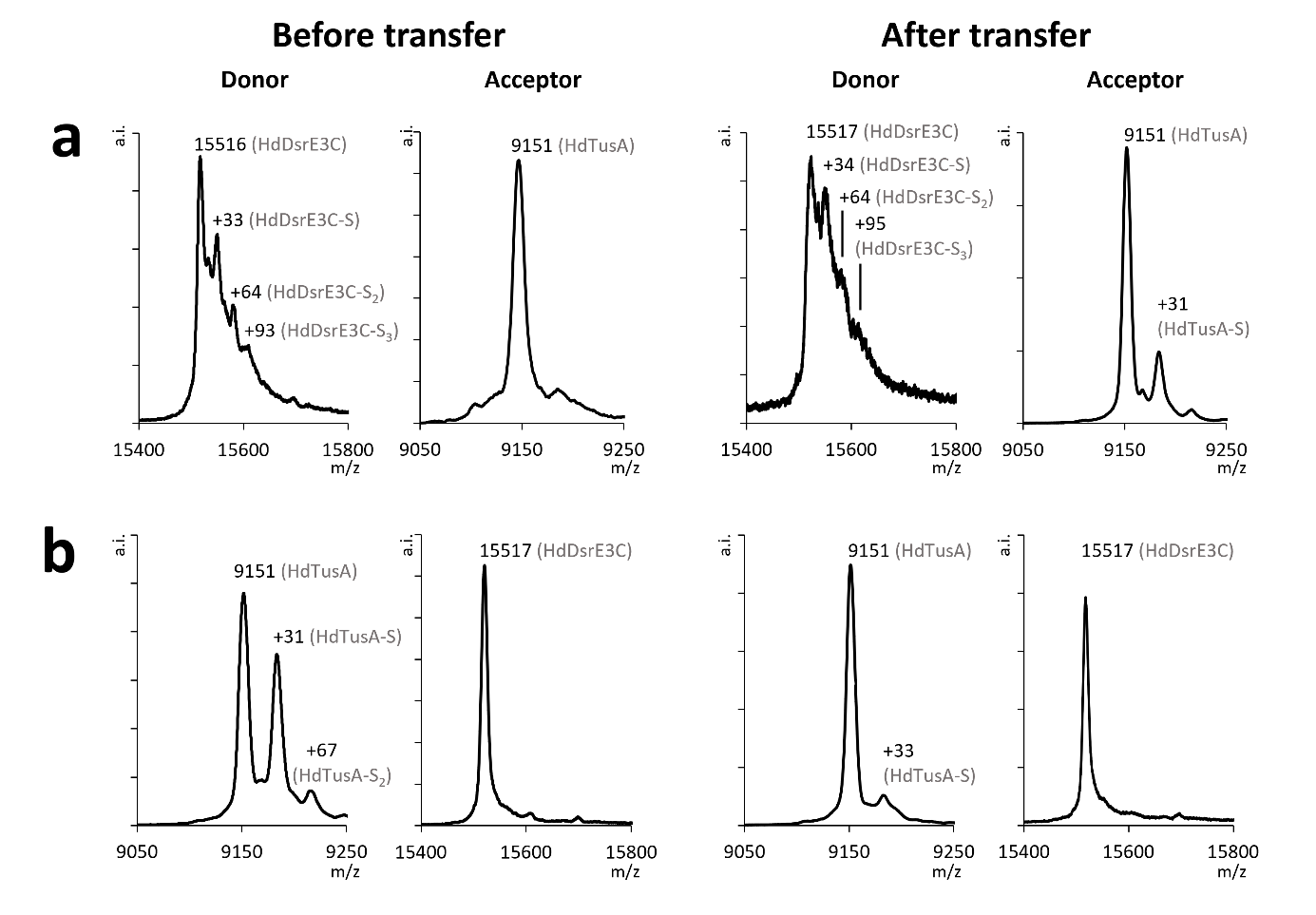

**FIGURE S2.** (a**)** The efficient transfer of sulfane sulfur from DsrE3C to TusA. (b) The transfer of sulfane sulfur from HdTusA to HdDsrE3C led to loss of sulfur from persulfurised HdTusA. The persulfuration of TusA was confirmed before mixing with HdDsrE3C (b donor before transfer), but the signal for this species was almost completely absent in the sample after incubation with HdDsrE3C (b donor after transfer). Neither a mass increase of 32 Da nor a change in ionization properties was observed for the HdDsrE3C after this reaction (b acceptor after transfer). Therefore, we conclude, that a sulfur transfer was not possible from HdTusA to HdDsrE3C.
