## Supplementary material for "A cascade of sulfur transferases delivers sulfur to the sulfur-oxidizing heterodisulfide reductase-like complex": Figures S1-S4, Tables S1-S7: Figure S3 Transfer_TkTusA_TkDsrE3B.docx

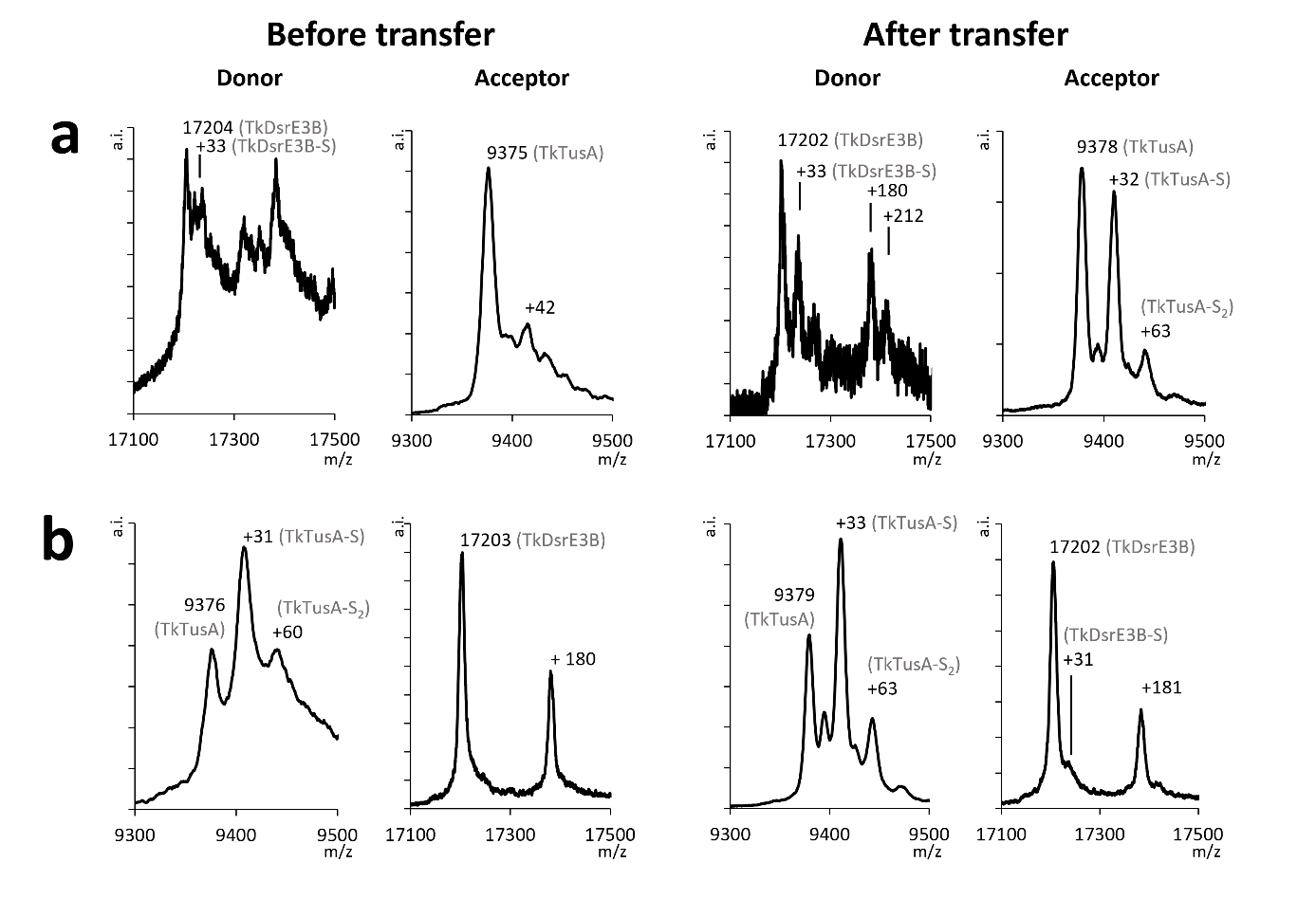

**FIGURE S3.** (a) The efficient transfer of sulfane sulfur from TkDsrE3B to TkTusA. Left panels: TkDsrE3B as persulfurated donor after treatment with polysulfide and unmodified reduced TkTusA as acceptor; right panels: TkDsrE3B (donor) and TkTusA (acceptor) after the transfer reaction. (b) Transfer of sulfane sulfur from TkTusA to TkDsrE3B is inefficient. Left panels: TkTusA as persulfurated donor after treatment with polysulfide and unmodified reduced TkDsrE3B as acceptor; right panels: TkTusA and TkDsrE3B after the transfer reaction.
