## Supplementary material for "A cascade of sulfur transferases delivers sulfur to the sulfur-oxidizing heterodisulfide reductase-like complex": Figures S1-S4, Tables S1-S7: Figure S4 Transfer_HdRhd.docx

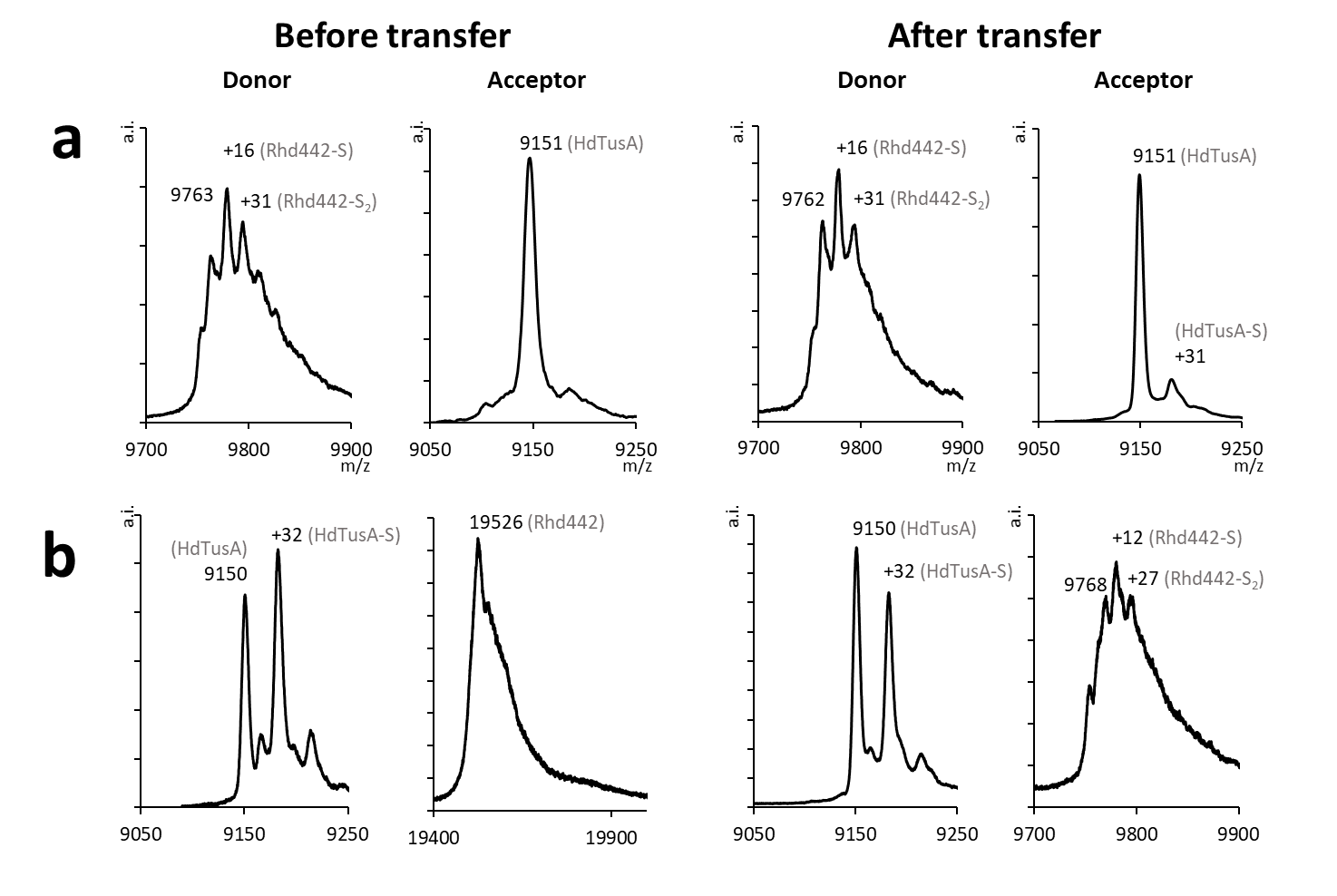

**FIGURE S4.** (a) Transfer of sulfane sulfur between HdRhd442 to HdTusA. Left panels: HdRhd442 as persulfurated donor after treatment with polysulfide and unmodified reduced HdTusA as acceptor. Right panels: HdRhd442 (donor) and HdTusA (acceptor) after the transfer reaction. (b) Left panels: HdTusA as persulfurated donor after treatment with polysulfide and unmodified reduced HdRhd442 as acceptor. Right panels: HdTusA (donor) and HdRhd442 (acceptor) after the transfer reaction.
