## Supplementary material for "A cascade of sulfur transferases delivers sulfur to the sulfur-oxidizing heterodisulfide reductase-like complex": Figures S1-S4, Tables S1-S7: Table S2 Sulfur loading TusA.docx

**TABLE S2.** Sulfur loading of TusA proteins from *H. denitrificans, Ts. sibirica,* *Thioalkalivibrio* sp. K90mix and *Aquifex aeolicus* with various inorganic sulfur compounds and oxidized glutathione (GSSG) detected by mass spectrometry**.** Numbers in parentheses represent mass increases. -, no modification; G, glutathione.

| **Protein** | **Expected masses [Da]** | **Detected masses [Da]** | **Modification** |
| --- | --- | --- | --- |
| HdTusA + S_2_O_3_^2-^ | 9149 | 9149 | - |
| HdTusA + S_4_O_6_^2-^ | 9149 | 9150 9148 (Δ34) 9212 (Δ62) 9262 (Δ112) 9296 (Δ146) | - -S -S_2_ -S_2_O_3_ -S_3_O_3_ |
| HdTusA + GSSG | 9149 | 9148 9453 (Δ304) | - -SG |
| HdTusA + Polysulfide | 9149 | 9149 9180 (Δ31) 9208 (Δ60) 9238 (Δ90) | - -S -S_2_ -S_3_ |
| Aq TusA + S_2_O_3_^2-^ | 9610 | 9611 | - |
| Aq TusA + GSH | 9610 | 9612 | - |
| Aq TusA + GSSH | 9610 | 9612 | - |
| Aq TusA + S_4_O_6_^2-^ | 9610 | 9610 9642 (Δ32) 9724 (Δ114) 9755 (Δ145) | - -S -S_2_O_3_ -S_3_O_3_ |
| Aq TusA + Polysulfide | 9610 | 9612 9643 (Δ31) | - -S |
| Aq TusA + Na_2_S | 9610 | 9611 9643 (Δ32) | - -S |
| TkTusA + S_2_O_3_^2-^ | 9376 | 9375 | - |
| TkTusA + S_4_O_6_^2-^ | 9376 | 9376 9410 (Δ32) 9490 (Δ111) 9525 (Δ146) 9549 (Δ171) | - -S -S_2_O_3_ -S_3_O_3_ -S_4_O_3_ |
| TkTusA + GSSG | 9376 | 9375 9680 (Δ306) | - -SG |
| TkTusA + Polysulfide | 9376 | 9380 9412 (Δ32) 9445 (Δ64) | None -S -S_2_ |
| TsTusA + S_2_O_3_^2-^ | 9393 | 9398 | - |
| TsTusA + S_4_O_6_^2-^ | 9393 | 9393 9426 (Δ32) 9505 (Δ112) | - -S -SO_3_ |
| TsTusA + GSSG | 9393 | 9392 | - |
| TsTusA + Polysulfide | 9393 | 9395 9427 (Δ32) 9459 (Δ64) | - -S -S_2_ |
