## Supplementary material for "A cascade of sulfur transferases delivers sulfur to the sulfur-oxidizing heterodisulfide reductase-like complex": Figures S1-S4, Tables S1-S7: Table S3 Sulfur transfer assays.docx

**Table S3. Detection of sulfur transfer between DsrE proteins, TusA and Rhd442.** Numbers in parentheses represent mass increases. -, no modification.

| **Donor** | | | |  | **Acceptor** | | | |
| --- | --- | --- | --- | --- | --- | --- | --- | --- |
| **Protein** | **Expected masses [Da]** | **Detected masses [Da]** | **Modifications** |  | **Protein** | **Expected masses [Da]** | **Detected masses [Da]** | **Modification** |
| HdTusA-S | 9149 + 32x | 9151 9184 (Δ32) | - -S | → | HdDsrE3C | 15520 + 32x? | 15517 | - |
| HdDsrE3C-S | 15520 + 32x | 15517  15551 (Δ34)  15581 (Δ64) 15612 (Δ95) | - -S -S_2_ -S_3_ | → | HdTusA | 9149 + 32x? | 9151 9182 (Δ32) | - -S |
| HdDsrE3C-S | 15520 + 32x? | 15519 15541 (Δ 32) | - -S | → | HdTusA Cys^13^Ser | 9133 + 32x? | 9135 | - |
| HdDsrE3C Cys^83^Ser -S | 15503 + 32x? | 15503 15535 (Δ32) 15567 (Δ64) | - -S -S_2_ | → | HdTusA | 9149 + 32x? | 9149 9181 (Δ32) | - -S |
| HdDsrE3C Cys^84^Ser -S | 15503 + 32x? | 15503 | - | → | HdTusA | 9149 + 32x? | 9148 | - |
| TkTusA-S | 9376 + 32x | 9379 9412 (Δ33) 9442 (Δ63) | - -S -S_2_ | → | TkDsrE3B | 17202 +32x? | 17202 17234 (Δ31) 17383 (Δ181) | - -S -glucose |
| TkDsrE3B-S | 17202 + 32x | 17202 17235 (Δ33) 17382 (Δ180) 17414 (Δ212) | - -S -glucose -S + -glucose | → | TkTusA | 9376 + 32x? | 9378 9410 (Δ32) 9441 (Δ63) | - -S -S_2_ |
| TsTusA-S | 9393 + 32x | 9396 9427 (Δ31) 9457 (Δ61) | - -S -S_2_ | → | TsDsrE3B | 16620 + 32x? | 16622 16653 (Δ31) 16801 (Δ179) | - -S -glucose |
| TsDsrE3B-S | 16620 + 32x | 16622 16653 (Δ31) 16684 (Δ62) 16720 (Δ98) | NM -S -S_2_ -S_3_ | → | TsTusA | 9393 +32x? | 9395 9427 (Δ32) | - -S |
| HdRhd442-S | 19526 + 32x | 19554 9762 (z = 2) 9778 (Δ16) (z = 2) 9793 (Δ31) (z = 2) | -S - -S -S_2_ | → | HdTusA | 9149 + 32x? | 9149 9181 (Δ32) | -S |
| HdRhd442-S | 19526 +32x | 19554 9763 (z = 2) 9779 (Δ16) (z = 2) 9795 (Δ32) (z = 2) 9810 (Δ47) (z = 2) | -S - -S -S_2_ -S_3_ | → | HdDsrE3C | 15520 + 32x? | 15519 15550 (Δ32) | -S |
