## Supplementary material for "A cascade of sulfur transferases delivers sulfur to the sulfur-oxidizing heterodisulfide reductase-like complex": Figures S1-S4, Tables S1-S7: Table S4.docx

Table S4: Proteins identified by mass spectrometry after re-purification of His-tagged AqTusA incubated with an *Aq. aeolicus* soluble extract (Band 1).

| **Accession** | **Description** | **Locus tag** | **MW** | **Coverage** | **Score** | **# PSMs** | **# Unique Peptides** |
| --- | --- | --- | --- | --- | --- | --- | --- |
|  | **TusA_tag** | aq_388a | 9,611 | **90** | 850 | **237** | **10** |
| O66901 | hydrogenase expression/formation protein B | aq_671 | 28,997 | 38 | 37 | 13 | 10 |
| O67481 | OsmC/Ohr family protein | aq_1515 | 15,448 | 46 | 16 | 6 | 5 |
| O67877 | Acetoin utilization protein | aq_2110 | 34,807 | 21 | 10 | 4 | 4 |
| O67476 | O-methyltransferase | aq_1507 | 24,296 | 17 | 9 | 4 | 4 |
| O66829 | Uncharacterized RNA pseudouridine synthase | aq_554 | 27,845 | 13 | 8 | 3 | 3 |
| **O66711** | **DsrE3C** | **aq_390** | 15,584 | **31** | 8 | **3** | **3** |
| **O66720** | **LbpA** | **aq_402** | 15,813 | **27** | 7 | **3** | **3** |
| O66599 | Rieske-I iron sulfur protein | aq_234 | 26,858 | 14 | 6 | 3 | 3 |
| O66859 | DUF2795 domain-containing protein | aq_600 | 18,31 | 17 | 6 | 3 | 3 |
| O67802 | Putative carboxymethylenebutenolidase | aq_1997 | 26,34 | 10 | 4 | 2 | 2 |
| O67045 | iron-sulfur cluster assembly scaffold protein IscU | aq_896 | 17,519 | 13 | 4 | 2 | 2 |
| O66523 | 30S ribosomal protein S16 | aq_123 | 13,03 | 17 | 4 | 2 | 2 |
| O67717 | CoA-binding domain-containing protein | aq_1869 | 15,326 | 15 | 4 | 2 | 2 |
| O66857 | DUF302 domain-containing protein | aq_598 | 14,51 | 19 | 3 | 2 | 2 |

Proteins from band 1 of the native gel (Figure 8 a) were analyzed.

Accession: accession number in UniProt database. MW: theoretical molecular weight of protein in kDa. Coverage: percent protein sequence coverage by the matching peptides. # Peptides: number of distinct peptides matching to protein sequence and unique to this protein. # PSMs: peptide spectrum match number (given by the algorithm corresponding to the total number of identified peptide sequences for the protein, including those redundantly identified). The *Aq. aeolicus* TusA protein (WP_024015099) is not referenced in Uniprot and the corresponding sequence was manually added in the database.
