## Supplementary material for "A cascade of sulfur transferases delivers sulfur to the sulfur-oxidizing heterodisulfide reductase-like complex": Figures S1-S4, Tables S1-S7: Table S5.docx

**Table S5**: Proteins, encoded by the *Aq. aeolicus* *hdr*-like cluster, identified by mass spectrometry after re-purification of His-tagged TusA incubated with an *Aq. aeolicus* soluble extract (second experiment).

|  |  |  |  |  | **Control** |  |  |  | **TusA** |  |  |
| --- | --- | --- | --- | --- | --- | --- | --- | --- | --- | --- | --- |
| **Accession** | **Description** | **Locus tag** | **MW** | **Coverage** | **Score** | **# Peptides** | **# PSMs** | **Coverage** | **Score** | **# Peptides** | **# PSMs** |
|  | TusA_tag | aq_388a | 9,611 | - | - | - | - | 88 | 861 | 13 | 211 |
| O66711 | DsrE3C | aq_390 | 15,584 | - | - | - | - | 46 | 91 | 6 | 28 |
| O66718 | heterodisulfide reductase subunit B HdrB2 | aq_400 | 37,221 | - | - | - | - | 42 | 68 | 14 | 20 |
| A0A193BL08 | heterodisulfide reductase subunit B HdrB1 | aq_392 | 51,938 | - | - | - | - | 32 | 46 | 12 | 15 |
| O66712 | heterodisulfide reductase subunit C HdrC1 | aq_391 | 29,438 | - | - | - | - | 35 | 19 | 7 | 7 |
| O66710 | DsrE2A | aq_389 | 19,88 | - | - | - | - | 13 | 7 | 2 | 2 |
| O66715 | heterodisulfide reductase subunit A HdrA | aq_395 | 38,517 | 27 | 25 | 8 | 9 | 67 | 83 | 18 | 25 |
| O66717 | heterodisulfide reductase subunit C HdrC2 | aq_398 | 29,823 | 7 | 7 | 3 | 3 | 20 | 16 | 5 | 6 |
| O66720 | LbpA2 | aq_402 | 15,813 | 13 | 7 | 2 | 2 | 32 | 20 | 3 | 5 |

“TusA” corresponds to the incubation of AqTusA with the soluble extract (second trial) and “Control” to the incubation of the extract without the AqTusA (negative control). Accession: accession number in UniProt database. MW: theoretical molecular weight of protein in kDa. Coverage: percent protein sequence coverage by the matching peptides. # Peptides: number of distinct peptides matching to protein sequence and unique to this protein. # PSMs: peptide spectrum match number (given by the algorithm corresponding to the total number of identified peptide sequences for the protein, including those redundantly identified). The *Aq. aeolicus* TusA protein (WP_024015099) is not referenced in Uniprot and the corresponding sequence was manually added in the database. HdrB1 was originally encoded by a pseudogene and corresponded to locus tags aq_392 and aq_394.^(1)^ “-“ means that the protein was not identified in the sample. The proteins AqsHdrH (aq_397) and AqDsrE3B (aq_401) were not identified in any sample in this experiment.
