## Supplementary material for "A cascade of sulfur transferases delivers sulfur to the sulfur-oxidizing heterodisulfide reductase-like complex": Figures S1-S4, Tables S1-S7: Table S6.docx

Table S6: Proteins identified by mass spectrometry after re-purification of His-tagged AqTusA incubated with an *Aq. aeolicus* soluble extract (bands 2 and 3).

| **Accession** | **Description** | **Locus tag** | **MW** | **Coverage** | **Score** | **# PSMs** | **# Unique Peptides** |
| --- | --- | --- | --- | --- | --- | --- | --- |
| **Band 2** |  |  |  |  |  |  |  |
| O66869 | Protease I | aq_618 | 18.961 | 71 | 321 | 104 | 13 |
| O67566 | Small ribosomal subunit protein uS8 | aq_1651 | 19.445 | 39 | 16 | 6 | 6 |
| O66780 | Thiol peroxidase | aq_488 | 18.561 | 31 | 11 | 5 | 5 |
| O67587 | ATP-dependent protease subunit HslV | aq_1671 | 19.276 | 38 | 10 | 5 | 5 |
| O66541 | Citrate synthase | aq_150 | 29.057 | 5 | 8 | 3 | 2 |
| **O66710** | **DsrE2A** | **aq_389** | **19.88** | **13** | **8** | **4** | **2** |
| O67527 | ATP synthase subunit delta | aq_1588 | 20.723 | 19 | 7 | 3 | 3 |
| O66981 | Cytochrome c552 CycB1 | aq_792 | 18.474 | 9 | 7 | 3 | 2 |
| O67903 | Peptidoglycan associated lipoprotein | aq_2147 | 22.842 | 16 | 7 | 3 | 3 |
| O67613 | dTTP/UTP pyrophosphatase | aq_1718 | 21.037 | 16 | 5 | 2 | 2 |
| O66616 | Uncharacterized protein | aq_255 | 18.849 | 8 | 5 | 2 | 2 |
| O66764 | Archaemetzincin | aq_459 | 20.324 | 11 | 5 | 2 | 2 |
| O67100 | Arginine biosynthesis bifunctional protein ArgJ | aq_970 | 41.717 | 6 | 4 | 2 | 2 |
| **O67573** | **LbpA3** | **aq_1657** | **18.215** | **19** | **4** | **2** | **2** |
| O67742 | Orotate phosphoribosyltransferase | aq_1907 | 20.297 | 14 | 4 | 2 | 2 |
| O67722 | Large ribosomal subunit protein uL13 | aq_1877 | 17.057 | 11 | 4 | 2 | 2 |
| **Band 3** |  |  |  |  |  |  |  |
| O67294 | Copper chaperone PCu(A)C | aq_1253 | 17.191 | 55 | 158 | 54 | 11 |
| **O66711** | **DsrE3C** | **aq_390** | **15.584** | **56** | **46** | **15** | **7** |
| O66523 | Small ribosomal subunit protein bS16 | Aq_123 | 13.03 | 39 | 44 | 17 | 8 |
| O67481 | OsmC family peroxiredoxin | aq_1515 | 15.448 | 49 | 40 | 14 | 6 |
| O66486 | Small ribosomal subunit protein uS13 | aq_074 | 14.331 | 28 | 26 | 9 | 6 |
| **O66720** | **LpbA2** | **aq_402** | **15.813** | **15** | **21** | **6** | **4** |
| O66537 | Uncharacterized protein | aq_142 | 16.311 | 38 | 16 | 6 | 5 |
| O67065 | Ferredoxin-1 | aq_919.1 | 10.755 | 41 | 15 | 5 | 3 |
| O66485 | Small ribosomal subunit protein uS11 | aq_073 | 13.518 | 10 | 13 | 5 | 2 |
| O67227 | Uncharacterized protein | aq_1163 | 14.885 | 32 | 11 | 5 | 4 |
| O67562 | Large ribosomal subunit protein uL30 | aq_1644 | 14.454 | 37 | 11 | 4 | 4 |
| O66541 | Citrate synthase | aq_150 | 29.057 | 14 | 11 | 5 | 3 |
| O66582 | KaiC domain-containing protein | aq_204 | 32.931 | 16 | 10 | 4 | 4 |
| O67534 | Rhodanese domain-containing protein | aq_1599 | 17.3 | 30 | 10 | 4 | 4 |
| O66705 | CRISPR type III-B/RAMP module-associated protein Cmr5 | aq_384 | 15.66 | 25 | 7 | 3 | 3 |
| O66901 | Hydrogenase expression/formation protein B | aq_671 | 28.997 | 16 | 7 | 3 | 3 |
| P0A466 | Large ribosomal subunit protein bL12 | aq_1937 | 13.551 | 9 | 7 | 3 | 2 |
| **O66719** | **DsrE3B** | **aq_401** | **14.887** | **28** | **7** | **3** | **2** |
| O66910 | Arsenate reductase ArsC | aq_685 | 16.233 | 20 | 6 | 3 | 3 |
| O67706 | Uncharacterized protein | aq_1854 | 17.702 | 11 | 6 | 2 | 2 |
| O67424 | Minor pilin | aq_1433 | 12.83 | 17 | 5 | 2 | 2 |
| O67548 | Metallo-beta-lactamase domain-containing protein | aq_1625 | 28.245 | 9 | 5 | 2 | 2 |
| O67570 | Large ribosomal subunit protein uL14 | aq_1654 | 13.263 | 25 | 5 | 2 | 2 |
| O67073 | NADPH-dependent 7-cyano-7-deazaguanine reductase | aq_931 | 15.321 | 18 | 5 | 2 | 2 |
| O67054 | Phosphoheptose isomerase | aq_908 | 20.693 | 16 | 4 | 2 | 2 |
| O66433 | Large ribosomal subunit protein uL23 | aq_012 | 12.363 | 23 | 5 | 2 | 2 |
| O67234 | Phosphohistidine phosphatase SixA | aq_1173 | 16.997 | 18 | 4 | 2 | 2 |
| O66862 | Acyl-[acyl-carrier-protein]--UDP-N-acetylglucosamine O-acyltransferase | aq_604 | 28.398 | 10 | 4 | 2 | 2 |

Proteins from bands 2 (about 18 kDa) and 3 (about 15 kDa) of the SDS gel (Figure 8b) were analyzed. Accession: accession number in UniProt database. MW: theoretical molecular weight of protein in kDa. Coverage: percent protein sequence coverage by the matching peptides. # Peptides: number of distinct peptides matching to protein sequence and unique to this protein. # PSMs: peptide spectrum match number (given by the algorithm corresponding to the total number of identified peptide sequences for the protein, including those redundantly identified). The *Aq. aeolicus* TusA protein (WP_024015099) is not referenced in Uniprot and the corresponding sequence was manually added in the database.
