## Supplementary material for "A cascade of sulfur transferases delivers sulfur to the sulfur-oxidizing heterodisulfide reductase-like complex": Figures S1-S4, Tables S1-S7: Table S7 Strains Primer and Plasmids.docx

**Table S1. Strains, plasmids and primers**

| **Strains primers or plasmids** | **Relevant genotype, description or sequence** | **Reference or source** |
| --- | --- | --- |
| **Strains** |  |  |
| *E. coli* 10-beta | Δ(ara-leu) 7697 araD139  fhuA ΔlacX74 galK16 galE15 e14-  ϕ80dlacZΔM15  recA1 relA1 endA1 nupG  rpsL (Str^R^) rph spoT1 Δ(mrr-hsdRMS-mcrBC) | New England Biolabs |
| *E. coli* DH5α | F^–^ φ80*lac*ZΔM15 Δ(*lac*ZYA-*arg*F)U169 *rec*A1 *end*A1 *hsd*R17(r_K_^–^, m_K_^+^) *pho*A *sup*E44 λ^–^*thi*-1 *gyr*A96 *rel*A1 | New England Biolabs |
| *E. coli* BL21 (DE3) | F^–^*omp*T *hsd*S_B_ (r_B_^–^, m_B_^–^) *gal dcm* (DE3) | Novagen |
| *Hyphomicrobium denitrificans* Δ*tsdA* | Sm^r^, in-frame deletion of *tsdA* in *H. denitrificans* Sm200 | (Koch & Dahl, 2018) |
| *Hyphomicrobium denitrificans* Δ*tsdA* Δ*dsrE3C* | Sm^R^, in-frame deletion of *dsrE3C* (Hden_0688) in *H. denitrificans* Δ*tsdA* | This work |
| *Hyphomicrobium denitrificans* Δ*tsdA* *dsrE-C83S* | Sm^R^, mutation of *dsrE3C* (Hden_0688) *C83 to S* in *H. denitrificans* Δ*tsdA* | This work |
| *Hyphomicrobium denitrificans* Δ*tsdA* *dsrE-C84S* | Sm^R^, mutation of *dsrE3C* (Hden_0688) *C83 to S* in *H. denitrificans* Δ*tsdA* | This work |
| **Primers** |  | This work |
| Hden0688 (*dsrE3C*)_Up_Fw_PstI | ATAT**CTGCAG**CCAATCTGCGTGGCGTTCCG | This work |
| Hden0688 (*dsrE3C*)_Up_Rev | CCCTGCCGTCCGAAAAATTCAATGCCACCTCCCCGATATG | This work |
| Hden0688 (*dsrE3C*)_Down_Fw | CATATCGGGGAGGTGGCATTGAATTTTTCGGACGGCAGGG | This work |
| Hden0688 (*dsrE3C*)_Down_Rev_XbaI | CATG**TCTAGA**TGCGCGTCGGTGATGCGATG | This work |
| Hden0688 (*dsrE3C*)_C83S_Fw | GTGAAATTTTTCTCCTGTTCTCCCAATCTC | This work |
| Hden0688 (*dsrE3C*)_C83S_Rev | GAGATTGGGAGAACAGGAGAAAAATTTCAC | This work |
| Hden0688 (*dsrE3C*)_C84S_Fw | GTGAAATTTTTCTGCTCTTCTCCCAATCTC | This work |
| Hden0688 (*dsrE3C*)_C84S_Rev | GAGATTGGGAGAAGAGCAGAAAAATTTCAC | This work |
| Hden0688 (*dsrE3C*)_NdeI_fw | CACG**CATATG**TTGGCCGAAAAACTTCTG | This work |
| Hden0688 (*dsrE3C*)_BamHI_rev | GCGT**GGATCC**TCAGTATGAAAGCACTTTG | This work |
|  |  | This work |
| TK90_0639 (*dsrE3B*)_NdeI_fw | AGAG**CATATG**ATGGCTGAACTGGGC | This work |
| TK90_0639_(*dsrE3B*)_BamHI_rev | CGA**GGATCC**AAAATTAAAACGTTATCGT | This work |
| ThisiDRAFT_1818 (*dsrE3B*)_NdeI_fw | AGA**CATATG**ACTGATGCAAGTC | This work |
| ThisiDRAFT_1818 (*dsrE3B*)_XhoI_rev | TTTTT**CTCGAG**TTAGAAGTTTATGAT | This work |
| HdenTusA_fw NdeI | CGACCA**CATATG**GCCGATCTGACAGTTGAT | This work |
| HdenTusA_rev_EcoRI | GCAAGC**GAATTC**TTATTTTTCGAACTGCGGGTGGCTCCAAGCGCTGGCCGCCGTGTGCTTGATCA | This work |
| Tk90TusA_fw_NdeI | AGACAC**CATATG**GCCAACTTTGACCAAGA | This work |
| TK90TusA_rev_EcoRI | CTTAGA**GAATTC**TTATTTTTCGAACTGCGGGTGGCTCCAAGCGCTGGACTTCTTCACGAGGAAGTAGAACTT | This work |
| ThisiTusA_fw_NdeI | ACACACG**CATATG**GCAAATTTTGACCTAGAACT | This work |
| ThisiTusA_rev_EcoRI | GCCAGT**GAATTC**TTATTTTTCGAAGTGCGGGTGGCTCCAAGCGCTGCTCTTGCGGATC | This work |
| HdenTusA_C13S_fw | GGCACGAACTCTCCTATCCCGATTTTGAAG | This work |
| HdenTusA_C13S_rev | CTTCAAAATCGGGATAGGAGAGTTCGTGCC | This work |
| Aq388a_NdeI_fw | TAGTT**CATATG**GCTACAATAACACCTGACAAGG | This work |
| Aq388a_XhoI_rev | AGT**CTCGAG**TCCTTTTTTCCTTATGTAGTAGATGTACTTACC | This work |
| AqT388a _C17_fw | CGATACTTCCGGACTTAACTCTCCTCTGCCCGTG | This work |
| Aq388a _C17_rev | CACGGGCAGAGGAGAGTTAAGTCCGGAAGTATCG | This work |
| Aq388a _C54_fw | GATATTCCAGCGTTCTCTCAAAGGACTGGACAC | This work |
| Aq388a _C54_rev | GTGTCCAGTCCTTTGAGAGAACGCTGGAATATC | This work |
| HdRhd442_fw_NdeI | CGAT**CATATG**AGTCAAGAAACCTGC | This work |
| HdRhd442_rev_HindIII | GATC**AAGCTT**CTCGTCGTCATCTTTCAGC | This work |
|  |  | This work |
|  |  | This work |
|  |  | This work |
|  |  | This work |
|  |  | This work |
|  |  | This work |
|  |  | This work |
|  |  | This work |
|  |  | This work |
|  |  | This work |
|  |  | This work |
|  |  | This work |
|  |  | This work |
|  |  | This work |
|  |  | This work |
|  |  | This work |
|  |  | This work |
| **Plasmids** |  |  |
| pET-22b(+) | Ap^r^ | Novagen |
| pET-15a(+) | Ap^r^ | Novagen |
| pET-15a-Hd-DsrE3C | Ap^r^, pET-15a with dsrE3C (Hden_0688) insertion between NdeI and BamHI | This work |
| pET-15a-Hd-DsrE3C-C83S | Ap^r^, pET-15a-Hd-DsrE3C with Cys^83^Ser exchange | This work |
| pET-15a-Hd-DsrE3C-C84S | Ap^r^, pET-15a-Hd-DsrE3C with Cys^84^Ser exchange | This work |
| pET-15a-TK90-DsrE3B | Ap^r^, pET-15a with dsrE3B (TK90_0639) insertion between NdeI and BamHI | This work |
| pET-15a-Ts-DsrE3B- | Ap^r^, pET-15a with dsrE3B (ThisiDRAFT_2311) insertion between NdeI and BamHI | This work |
| pET22b-HdenTusA | Ap^r^, pET-22b with tusA (Hden_0698) insertion between NdeI and BamHI | This work |
| pET22b-HdenTusA-C13S | Ap^r^, pET22b-HdenTusA with Cys^13^Ser exchange | This work |
| pET22b-TK90TusA | Ap^r^, pET-22b with tusA insertion between NdeI and BamHI | This work |
| pET22b-ThisiTusA | Ap^r^, pET-22b with tusA insertion between NdeI and XhoI | This work |
|  | Ap^r^, pET-22b-SoxR-N-Strep with Cys^50^Ser ans Cys^116^Ser exchanges | This work |
| pk18*mobsacB*-Tc | Km^r^, Tc^r^ pHP45ΩTc tetracycline cassette inserted into pk18*mobsacB* using SmaI | (Li et al., 2022) |
| pk18*mobsacB*-Tc | Km^r^, Tc^r^ pHP45ΩTc tetracycline cassette inserted into pk18*mobsacB* using SmaI | (Li et al., 2022) |
| pk18*mobsacB*_Tc_dsrE3C-C83S | Km^r^, Tc^r^, 2.08 kb SOE PCR fragment implementing mutation C83S of *dsrE3C* cloned into pk18mobsacB-Tc using PstI and XbaI restriction sites | (Li et al., 2022) |
| pk18*mobsacB*_Tc_dsrE3C-C84S | Km^r^, Tc^r^, 2.08 kb SOE PCR fragment implementing mutation C84S of *dsrE3C* cloned into pk18mobsacB-Tc using PstI and XbaI restriction sites | (Li et al., 2022) |
| pk18*mobsacB*_Tc_Δ*dsrE* (Hden0688) | Km^r^, Tc^r^, 2.08 kb SOE PCR fragment implementing deletion of *dsrE3C* cloned into pk18mobsacB-Tc using PstI and XbaI restriction sites | This work |
